# *fruitless* functions downstream of *doublesex* to promote sexual dimorphism of the gonad stem cell niche

**DOI:** 10.1101/454546

**Authors:** Hong Zhou, Cale Whitworth, Caitlin Pozmanter, Megan C. Neville, Mark Van Doren

## Abstract

**Background:** *doublesex* (*dsx*) and *fruitless* (*fru*) are the two downstream transcription factors that actuate *Drosophila* sex determination. While *dsx* assists *fru* to regulate sex-specific behavior, whether *fru* collaborates with *dsx* in regulating other aspects of sexual dimorphism remains unknown. One important aspect of sexual dimorphism is found in the gonad stem cell (GSC) niches, where male and female GSCs are regulated to create large numbers of sperm and eggs.

**Results:** Here we report that Fru is expressed male-specifically in the GSC niche and plays important roles in the development and maintenance of these cells. Unlike previously studied regulation of sex-specific Fru expression, which is regulated by alternative splicing by Transformer (Tra), we show that male-specific expression of *fru* is regulated downstream of *dsx*, and is independent of Tra. Regulation of *fru* by *dsx* also occurs in the nervous system. *fru* genetically interacts with *dsx* to support maintenance of the hub throughout development. Ectopic expression of *fru* inhibited female niche formation and partially masculinized the ovary. *fru* is also required autonomously for cyst stem cell maintenance and cyst cell survival. Finally, we identified a conserved Dsx binding site upstream of *fru* promoter P4 that regulates *fru* expression in the hub, indicating that *fru* is likely a direct target for transcriptional regulation by Dsx.

**Conclusions:** These findings demonstrate that *fru* acts outside the nervous system to influence sexual dimorphism and reveal a new mechanism for regulating sex-specific expression of *fru* that is regulated at the transcriptional level by Dsx, rather than by alternative splicing by Tra.

## INTRODUCTION

In sexually reproducing animals, the proper production of gametes and successful copulation are equally critical for reproductive success. It is therefore important that both the gonad and the brain know their sexual identity. The Doublesex/Mab-3 Related Transcription Factors (DMRTs) act downstream of sex determination and play an evolutionarily conserved role to establish and maintain sexual dimorphism in the gonad [1]. Meanwhile, sexual dimorphism in other tissues such as the brain is controlled, to varying degrees in different animals, through autonomous control by the sex determination and non-autonomous signaling from the gonads[2, 3]. In many invertebrate species, another sex-determination gene *fruitless* (*fru*), which encodes several BTB-Zinc finger transcription factors, plays a central role in controlling sexual orientation and courtship behaviors [4]. How sex determination in the gonad and the nervous system is related and coordinated in these species remains unclear.

The founding member of the DMRT family is *Drosophila doublesex* (*dsx*). *dsx* and *fru* are alternatively spliced by the sex determination factor Transformer (Tra) to produce sex-specific protein isoforms. It was once thought that *dsx* controls sexual dimorphism outside the nervous system while *fru* regulates sex-specific nervous system development and behavior. But growing evidence shows that *dsx* is also required to specify the sex-specific neural circuitry and regulate courtship behaviors [5, 6] and cooperates with *fru* [7–10]. However, whether *fru* acts along with *dsx* to control sexual dimorphism outside the nervous system remains unknown.

The *fru* gene locus is a complex transcription unit with multiple promoters and alternative splice forms (Figure 1A). Sex-specific regulation of *fru* was only known to occur through alternative splicing of transcripts produced from the P1 promoter by Tra [11, 12]. The downstream promoters (P2-P4) produce Fru isoforms (collectively named as Fru^Com^) encoded by transcripts that are common to both sexes and are required for viability in both males and females. *fru* P1 transcripts have only been detected in the nervous system [13], suggesting that sex-specific functions of *fru* are limited to neural tissues. However, Fru^Com^ is expressed in several non-neural tissues, including sex-specific cell types of the reproductive system [13, 14]. Further, from a recent genome-wide search for putative DSX targets, we identified *fru* as a candidate gene that is regulated by Dsx ([15] and Fig S5). These data raise the possibility that *fru* functions cooperatively with *dsx* to regulate gonad development.

**Figure 1.**
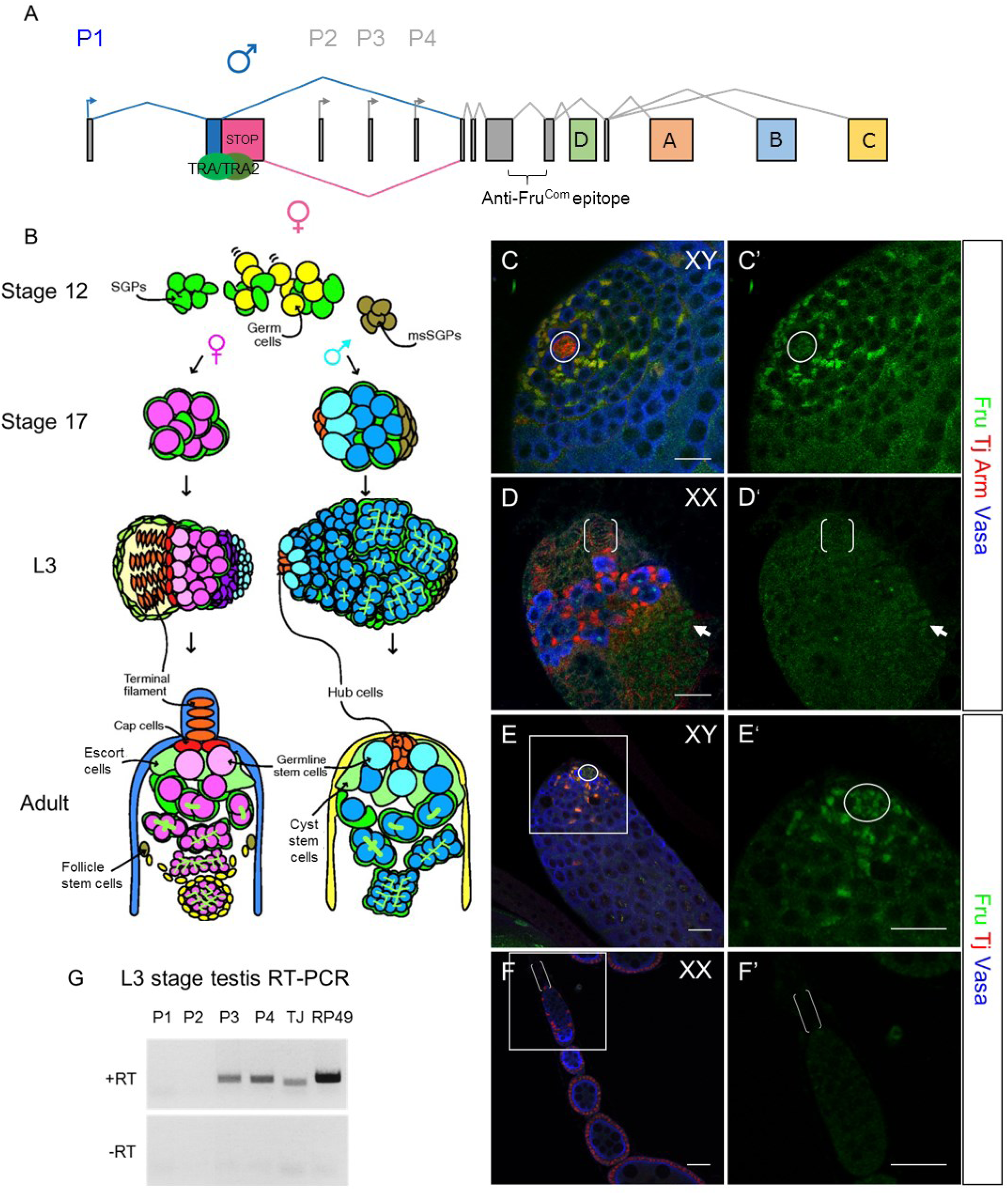
**Fruitless is expressed male-specifically in the germline stem cell niche and is independent of Fru^M^.** (A) Schematic of the *fruitless* (*fru*) gene locus. (B) Development of the female and male germline stem cell (GSC) niche. (C-D) late L3 stage larval gonads. The arrow in (D) indicates weak Fru expression in terminal epithelial cells. (E-F) Adult testis and ovary. Fru expression in the GSC niche is shown at a higher magnitude in (E’ and F’). Scale bars represent 20 µm. Circles: hubs; brackets: TFs. (G) RT-PCR of late L3 stage testes with promoter-specific primers. *tj* and *rp49* primers were used as positive controls.

The stem stem cell niche is a key component of the gonad that provides signals to the germline stem cells (GSCs) to regulate gametogenesis [16]. Sexual differences within the adult GSC niche have been well-characterized [16]. Critical components of the niche are the hub in males and terminal filaments (TFs) and cap cells in females (Figure 1B). Other important cell types include cyst stem cells (CySCs) and cyst cells in males and escort cells, follicle stem cells (FSCs) and follicle cells in females. The hub is a tight cluster of postmitotic cells that form during the last stages of embryogenesis [17]. In contrast, female niche specification starts in late 3^rd^ larval instar when stacks of terminal filament cells are specified from cells forming the apical cap, and completes at the larval-pupal transition with TFs inducing the specification of cap cells from intermingle cells [18–20]. Recently, it was shown that one important role *dsx* plays is to maintain the hub fate in the 3^rd^ instar larval (L3) stage and to prevent sex reversal (Camara *et al.*, submitted). In the absence of *dsx*, both XX and XY gonads initially follow the male path to form a hub in the embryonic stage, but later undergo stochastic sexual-fate reprogramming in the L3 stage. Eventually, half of both XX and XY gonads form TFs in place of the hub, while the other half maintain the hub. The genes and pathways that function downstream of *dsx* to regulate male vs. female gonad niche fate remain elusive.

To test if *dsx* and *fru* act in concert to regulate sexual development of the gonad, we investigated *fru* expression and function in the gonad. We found that Fru is expressed male-specifically in the GSC niche and functions downstream of *dsx* to regulate the development and maintenance of the male GSC niche. Our analyses show that *fru* is required in *dsx* mutant gonads to prevent hub-to-TF fate conversion and is sufficient to masculinize the developing female GSC niche. Fru also functions in the cyst stem cell (CySC) lineage to maintain CySC fate and to regulate spermatogonia cyst survival. Finally, we showed that *fru* P4 promoter is directly regulated by Dsx, through at least one evolutionarily conserved Dsx binding site. These results provide new insights into the organization of the *Drosophila* sex determination pathway and how the downstream regulators Dsx and Fru cooperate to control sexual dimorphism in the gonad and brain.

## RESULTS

### Male-specific Fruitless expression in the testis

To examine Fru expression in the gonad, we used the anti-Fru^Com^ antibody that recognizes all Fru isoforms [14]. Interestingly, we found that Fru has a dynamic and male-specific pattern of expression within the developing gonad. While the gonad forms during embryogenesis and the hub and cyst stem cells are specified in the late embryo and early L1 stage [17, 21], no anti-FRU immunoreactivity was observed in the gonads of either sex at this time (Figure S1A-S1B’). Male-specific Fru expression was first observed in some late L2 stage gonads (Figure S1C-S1D’) but was only consistently observed in L3 stage gonads (Figure S1E-S1F’). In the L3 stage, we observed Fru immunoreactivity in the hub cells through co-immunostaining with the hub marker Armadillo (Arm) (Figure 1C). We also observed Fru expression in cyst stem cells and early cyst cells that express the lineage marker Traffic jam (Tj) [22]. Within the ovary, we did not observe Fru expression in the apical cap from which the terminal filaments will form, or in the Tj-positive somatic cells that are intermingled with the germ cells at this stage (Figure 1D). Occasionally, we detected weak Fru signal in the basal epithelium of the ovary. We did not observe Fru expression in the germ cells (Vasa-positive) of either sex. This male-specific expression pattern is maintained in the adult GSC niche. In the testis, we observed Fru colocalizing with Tj-expressing hub cells, cyst stem cells and early cyst cells (Figure 1E). In contrast, Fru is not expressed in the terminal filament cells or the Tj-expressing somatic cells of the germarium (Figure 1F).

Tra-mediated alternative splicing of P1 transcripts is the only mechanism that is known to generate male-specific Fru expression pattern. However, P1 expression was not detected in the male reproductive system by northern blot [13]. To test if Fru proteins detected by the anti-Fru^Com^ antibody were from the P1 transcript, we utilized an engineered *fru* allele, *fru*^*F*^, which generates female-spliced transcripts from P1 in both sexes. These transcripts do not encode functional Fru protein and lack the anti-Fru^Com^ antibody epitope [23]. We reasoned that if male-specific Fru expression in the gonad is due to sex-specific splicing of P1, the anti-Fru^Com^ immunoreactivity would be abolished in the *fru*^*F*^ mutant testes. However, we observed normal Fru expression in *fru*^*F*^ mutant adult testes, suggesting that Fru^M^ is not responsible for male-specific Fru expression (Figure S1G). Consistent with this, flies containing a modified *fru* locus expressing *Gal4* in place of the P1 transcripts (*fru*^*Gal4*^, [24]) did not exhibit any Gal 4 activity in the testis tip when combined with a *UAS-mCD8GFP* reporter (Figure S1H). To determine which promoter drives *fru* expression in the male GSC niche, we generated cDNA from L3 stage testes that lack innervation by the *fru*^*M*^-expressing neurons [25]. RT-PCR conducted with promoter-specific primers showed that transcripts generated from the P3 and P4 promoters were expressed whereas P1 and P2 transcripts are absent in the gonad (Figure 1G).

We conclude that Fru is expressed sex-specifically in the male somatic gonad, specifically in the region of the gonad stem cell niche, and that this expression is independent of the P1 promoter that is the only known promoter subject to sex-specific alternative splicing.

### Male-specific Fru expression is dependent on *dsx* and independent of alternative splicing by Tra

As the *fru* P1 transcript is the only transcript thought to be subject to alternative splicing by Tra, and our genomic analysis indicated that *fru* is a candidate Dsx target gene [15], we considered the possibility that sex-specific Fru expression in the gonad is regulated at the transcriptional level by Dsx. Normally, Tra acts to splice both *dsx* and *fru P1* into their female-specific isoforms. To test whether male-specific Fru expression is dependent on *dsx* instead of *tra*, we utilized a genetic background that expresses the active (female) form of *tra* but the male form of *dsx.* This test utilizes an allele of *dsx* that can only produce the male isoform, even in XX animals (XX; *dsx*^*D*^/*dsx*^*0*^, Figure 2A). In this test, if the sex-specific expression of Fru in the gonad is dependent on female-specific splicing by Tra, or other components of the sex determination cascade upstream of *dsx*, we would expect Fru to be regulated in the “female mode” and not be expressed in the gonad. In contrast, if Fru expression is regulated by Dsx, we would expect Fru to be expressed in the “male mode” in the stem cell niche similar to wild-type testes. In XX; *dsx*^*D*^/*dsx*^*0*^ animals, we found that the somatic gonad was fully masculinized, resulting in the formation of a male niche (Figure 2C, [26]). Further, we observed robust and consistent Fru expression in L3 stage gonads, which overlapped with Fas-3 and Tj in the hub cells and the early CySC lineage, and was indistinguishable from the XY siblings (Figure 2B-2C’). This result indicates that Fru expression in the gonad is dependent on *dsx* and independent of *tra*.

**Figure 2.**
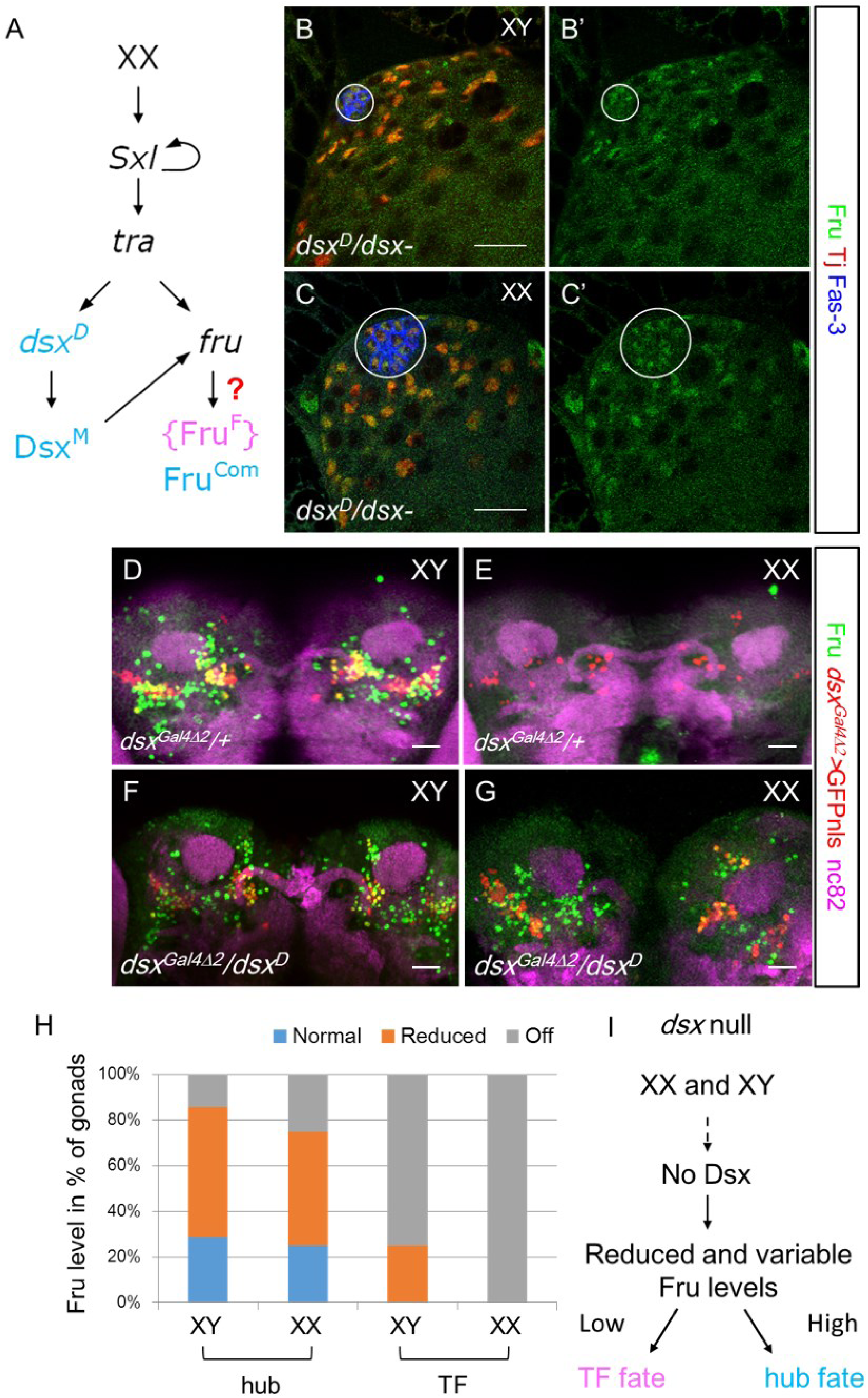
**Dsx is necessary and sufficient for sex-specific Fru^Com^ expression.** (A) Schematic of the experiment setup for (B-G). (B-C) Late L3 stage XX and XY gonads. Circle denotes the hub. Scale bars represent 20 µm. (D-G) Adult CNS immunostained with anti-Fru^Com^ antibody. Genotype as indicated. Scale bars represent 50 µm. (H) Distribution of Fru expression level detected by anti-Fru^Com^ antibody in *dsx*^3^/dsx^1^ gonads of the late L3 stage. Sample sizes are 4, 8, 12 and 7. (I) A model summarizing the Fru expression level in *dsx* null gonads.

Fru is male-specifically expressed in the CNS in a manner that is dependent on alternative splicing by Tra [11]. In adult males, Fru is expressed in a subset of sexually-dimorphic neurons of the posterior male brain, and partially overlaps with Dsx expression, which is indicated by *dsx*^*Gal4*Δ*2*^>GFP [27] (Figure 2D). In contrast, Fru is not expressed in the adult female CNS [14] (Figure 2E). We next sought to test if *dsx* also regulates sex-specific Fru expression in the CNS. Using the same genetic approach, where Tra is activated in the female mode while Dsx is expressed in the male mode (*XX; dsx*^*D*^/*dsx*^*0*^), we observed Fru expression similar to that observed in males. Fru was expressed in the posterior brain in a pattern similar to its expression in XY siblings and wild-type males (Figure 2D, 2F, and 2G). This result suggests that Fru expression in the CNS is dependent on Dsx in addition to the previously reported Fru expression that is regulated by sex-specific alternative splicing by Tra.

We then wanted to determine the expression of Fru in the gonad in the absence of *dsx* function. Based on studies from few known cases of Dsx targets [28–31], it is thought that DsxF and DsxM bind to the same target genes and regulate gene expression in the opposite directions. Therefore, we predicted that DsxM activates Fru expression in the testis while DsxF represses Fru expression in the ovary and that loss of *dsx* would cause Fru to be expressed at an intermediate level in both XX and XY gonads. In *dsx* mutants, half of both XX and XY gonads remain as hubs, while the other half switch to form TFs during the L3 stage. As a result, either a hub or TFs can be found in both XX and XY gonads (Camara *et al.,* submitted). We examined *dsx* null gonads at the late L3 stage and categorized the results by chromosomal sex and niche fate (hub vs. TFs, Figure 2H). Indeed, we found that *dsx* mutant gonad expressed Fru at an intermediate level, but that this level was highly variable (Figure S2A-S2D’). Further, the level of Fru expression correlated with whether the gonads had male or female niche structures; gonads with TFs were less likely to express Fru in the apical cap and TFs, while gonads with hubs tended to have higher levels of Fru expression.

Taken together, these findings indicate that sex-specific Fru expression in the gonad is regulated by *dsx*, and DsxM is required for robust and consistent Fru expression in the male niche while DsxF is required to repress Fru expression in the female niche. Further, the level of Fru expression in *dsx* mutants correlated with whether the gonad develops a male or female niche (Figure 2I). While we don’t know what regulates the variable level of Fru expression in the absence of *dsx*, this correlation suggests that *fru* influences male niche identity.

### *fru* functions downstream of *dsx* to maintain the male niche during development

The fact that some *dsx* mutant gonads switch from having hubs to TF during the L3 stage, at the time that the female niche normally develops, indicates that *dsx* is normally required in male gonads to maintain the male fate (Camara *et al.*, submitted). Fru is not expressed in the testis at the time of male niche formation during embryogenesis, but Fru expression initiates at the L2/L3 stage at the time that male niches must maintain hub fate, suggesting that Fru may be important for hub maintenance. We reasoned that if a higher Fru expression level is needed in *dsx* mutant gonads to maintain the hub identity or prevent TF formation, decreasing Fru dosage by removing one copy of *fru* would be sufficient to “tip the balance” and cause more gonads to switch to the formation of TFs. Conversely, if Fru expression is only a consequence of male-specific cell fate, changing Fru expression level would not alter the chances of a *dsx* mutant gonad developing a hub or TFs. *dsx* mutant gonads had a roughly equal chance of forming hubs or TFs (with another fraction forming no discernable niche structure, Figure 3A). When one copy of a *fru* allele was introduced as a heterozygote into this genetic background (*dsx*^*1*^/*dsx*^*3*^, *fru*^*Sat15*^/+), we observed that the fraction of XY gonads with hubs decreased while the fraction that formed TFs increased (Figure 3A). XX animals showed a similar bias towards the TF fate. These results suggest that, in *dsx* mutants, *fru* is required to maintain the hub fate and inhibit the TF fate.

**Figure 3.**
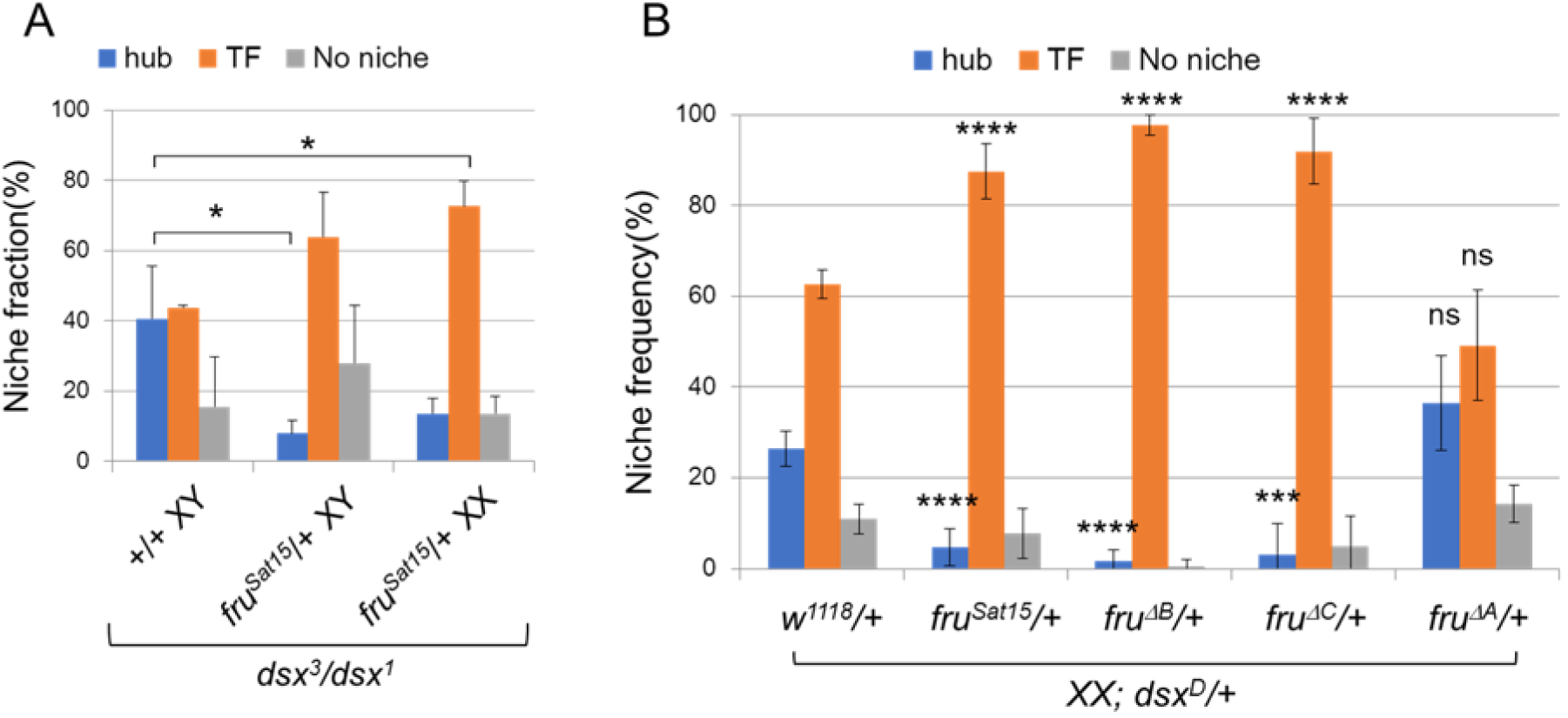
***fru* genetically interacts with *dsx* to maintain the male niche identity of *dsx* mutant gonads.** (A) Quantification of niche identity in 1-2 days old *dsx*^*3*^/*dsx*^*1*^ flies. (B) Quantification of niche identity in 1-2 days old XX *dsx*^*D*^/+ flies in the control background or with *fru* alleles. Data are presented as Mean ± SEM. Refer to Table S1 for sample sizes. Student’s t-test.

Fru proteins produced from the non-P1 promoters may contain one of four alternative ZnF domains (A, B, C, or D) that locate at the C-termini [32]. These Fru isoforms have distinct DNA binding motifs and play isoform-specific roles in the CNS [32]. Our RT-PCR result showed that all the exons are expressed in the larval testis (Figure S3A-S3B). To determine which *fru* C-terminal isoform is involved in promoting the hub fate, we switched to the *dsx*^*D*^/+ genetic background, where XX individuals simultaneously express DsxM and DsxF. The male and female isoforms of Dsx interfere with one another and these animals develop similar to *dsx* null animals [15]. In XX, *dsx*^*D*^/+ animals, we observed 62.73% of gonads formed TFs, 26.36% retained the hub and 10.91% had no identifiable niche (n=110). When one copy of a *fru* null allele was introduced into this background (XX; *dsx*^*D*^/*fru*^*Sat15*^), the fraction of gonads with TFs increased to 87.5% and the fraction of gonads with the hub decreased to 4.69% (n=256) (Figure 3B). This result confirmed the finding that reducing Fru dosage in *dsx* mutants causes more gonads to undergo hub-to-TF fate conversion. When genetic interaction assays were performed between *dsx*^*D*^ and isoform-specific *fru* mutant alleles, we found that *dsx*^*D*^/*fru*^Δ*B*^ and *dsx*^*D*^/*fru*^Δ*C*^ XX gonads had a similarly high frequency of TF fate whereas *fru*^Δ*A*^ did not affect the formation of hubs vs. TFs (Figure 3B and Table S1). From these results, we conclude that Fru^B^ and Fru^C^, but not Fru^A^, function to promote hub maintenance in *dsx* mutants.

### Loss of *fru* is not sufficient to cause gonad sex reversal

We next wanted to know whether loss of *fru* alone could cause gonad sex reversal. *fru* null, *fru*^Δ*B*^, and *fru*^Δ*C*^ mutant flies die in pupal stages [32], soon after the L3 stage when male niche fate must be maintained. We observe no morphological defect in the hub prior to lethality (Figure S4 A-D and data not shown), suggesting that loss of *fru* alone was not sufficient to cause a failure of hub maintenance. To determine whether *fru* helps to maintain the male niche in adult testes, we performed cell-type specific RNA-interference (RNAi) mediated knockdown of *fru*. Knockdown of *fru* in the hub using the *upd-Gal4* driver did not yield a hub phenotype (Figure S4E-S4F’). Knockdown of *fru* in the CySC lineage using the *tj-Gal4* also did not cause these cells to take on female morphology (Figure S4G and S4G’). From these results, we conclude that loss of Fru activity is not sufficient to cause testis sex reversal. It is worth noting, that when testes were examined 2 weeks after eclosure, we did observe an expansion of Tj-positive cyst cells in *tj>fru RNAi* testes compared to *tj>control RNAi* testes (Figure S4H-S4J), suggesting that *fru* has functions in regulating CySC lineage differentiation. However, since we observed no switching from hub to TF fate in *fru* mutants, we conclude that *dsx* regulates other targets in addition to *fru* to promote hub maintenance.

### *fru* is cell-autonomously required for cyst stem cell maintenance

To investigate further *fru* function in the CySC lineage, we generated GFP-positive *fru*-mutant clones using the MARCM technique [33] and asked if CySC clones could be generated and maintained. Marked control (*FRT82B*) CySC clones were observed in 67% (n=61), 56% (n=129) and 43% (n=56) of the testes examined at 2, 5, and 10 days post clone induction (pci), respectively (Figure 4A and Table S2). In contrast, CySC clones homozygously mutant for *fru*^*Sat15*^ were observed less frequently at 2 days pci (26%, n=46), lost rapidly by 5 days pci (1.8%, n=113), and were completely absent by 10 days pci (0%, n=78). *fru*^Δ*B*^ mutant CySCs were also observed at a low frequency at 2 days pci (29%, n=55), and were lost at a similar rate as *fru*^*Sat15*^ clones (5 days pci:4%, n=101; 10 days pci: 3% n=66). Interestingly, at 2 days pci, *fru*^Δ*C*^ CySC clones were observed in 57% of examined testes, similar to that of the control. The percentage of testes with *fru*^Δ*C*^ CySC clones decreased slightly at 5 days pci (30%, n=24). But by 10 days pci, only 5% of examined testes maintained *fru*^Δ*C*^ CySC clones (n=44), similar to the rate of *fru*^Δ*B*^ CySCs (Figure 4A). These results indicate that both Fru^B^ and Fru^C^ are required autonomously for CySC maintenance in adult testes and that Fru^B^ plays a more important role than FruC in the CySCs. Interestingly, we found that *fru*^Δ*B*^ mutant testes exhibited a significant reduction in anti-Fru^Com^ immunoreactivity whereas the signal in *fru*^Δ*C*^ mutant testes was comparable to the control testes (Figure S3C-S3F’).

**Figure 4.**
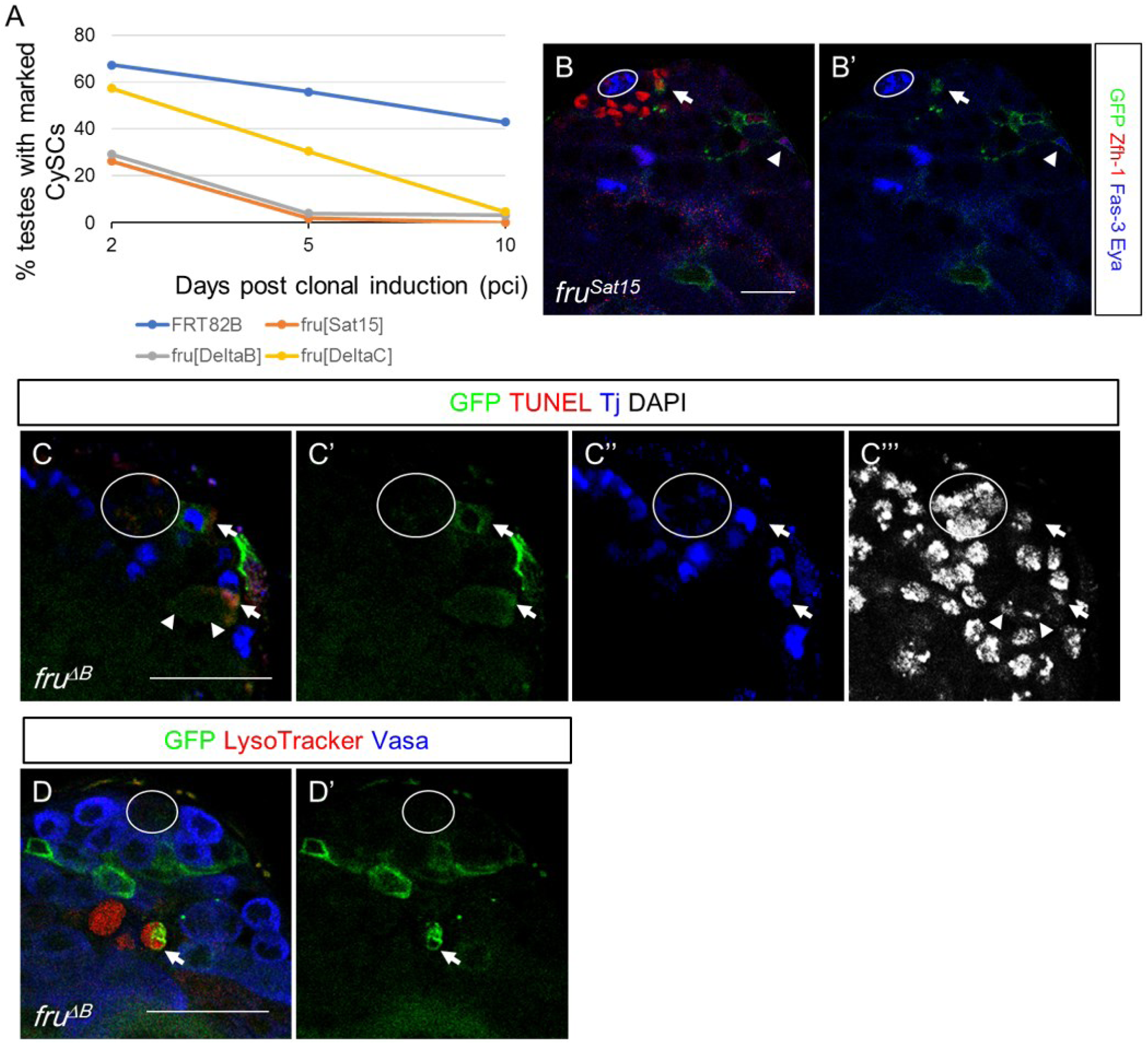
***fru* is cell-autonomously required for cyst stem cell maintenance and cyst cell survival.** (A) The percentage of control (*FRT80B*) and *fru* mutant (*fru*^*Sat15*^, *fru*^Δ*B*^, and *fru*^Δ*C*^) CySC clones maintained at the niche post clonal induction (pci). (B) A representative image at 4 days pci showing Zfh-1 and Eya expression in *fru*^*Sat15*^ CySC and cyst cell clones. Arrow denotes CySC clone and arrowhead denotes cyst cell clone. (C) A representative image at 4 days pci showing *fru*^Δ*B*^ cyst cell rather than CySC was positive for TUNEL. Arrow denotes *fru*^Δ*B*^ clones; arrowhead denotes germ cells encapsulated by the dying *fru*^Δ*B*^ cyst cell clones with diminished DAPI staining. (D) A representative image at 4 days pci showing a spermatogonia cyst encased by a *fru*^Δ*B*^ cyst cell clone (indicated by the arrow) was negative for Vasa and positive for LysoTracker. Circle denotes the hub. Scale bars represent 20 µm.

We next investigated the molecular mechanism by which *fru* regulates the CySC lineage. Two possible explanations of CySC loss are precocious differentiation and CySC cell death. Zfh-1 is expressed in CySCs and early differentiating cyst cells, while Eyes absent (Eya) is only expressed in later stages of cyst cell differentiation. In *fru* mutant clones at 2-4 days pci, the somatic cells closest to the hub still expressed Zfh-1 and did not express Eya, indicating they were not prematurely differentiating (Figure 4B). Similarly, *fru* mutant CySC did not exhibit signs of DNA fragmentation characteristic of apoptosis (TUNEL assay, Figure 4C), indicating that they also were not undergoing cell death. However, somatic cells further from the hub were more likely to be TUNEL positive when mutant for *fru* compared to wild-type controls. Further, germ cells encapsulated by *fru* mutant cyst cells were also more likely to exhibit condensed and reduced DAPI staining, suggesting that they were also dying. Such germ cells were also more likely to be positive for staining with LysoTracker, which is a characteristic of the Caspase-independent cell death exhibited by germ cells (Figure 4D). These results indicate that *fru* is required for CySC maintenance and for cyst stem cell viability and, likely as a consequence, the survival of spermatogenic cysts.

### Ectopic expression of Fru inhibits terminal filament formation and partially masculinizes the female niche

Though *fru* was not necessary for hub maintenance, we next asked whether *fru* was sufficient to cause defects in normal female niche development. We expressed the Fru^B^ (*UAS-fruB*) isoform in *dsx-*expressing cells of the developing ovary using *dsx-Gal4* [27, 34]. Engrailed (En) is a TF-specific marker and is required for specification of TF cells from the apical cap [35]. When white prepupae (WPP) were examined, control ovaries lacking the *UAS* transgene all had groups of 6-8 disc-shaped cells expressing En aligning at the base of the apical cap (n=7) (Figure 5A). In contrast, ovaries expressing Fru^B^ failed to robustly express En or intercalate En-expressing cells into filaments (n=25) (Figure 5B). To determine if Fru^B^ overexpression masculinized the female niche, we examined the male-specific niche marker, Escargot (Esg), with an enhancer trap (*esg*^*M5-4*^) that reports *esg* activity through the expression of LacZ [17, 36]. We observed strong expression of esg-LacZ in the hub of control testes (n=6), and no lacZ expression throughout the control ovary (Figure 5C-5D). In the WPP stage ovaries ectopically expressing Fru^B^, we detected a high level of esg-LacZ expression in the apical cap region (n=23) (Figure 5E). However, we did not observe any evidence for the formation of hubs in these gonads. Proteins produced from the *fru* P1 transcript in males (Fru^M^) have an N-terminal domain not found in proteins from other transcripts. Interestingly, ectopic expression of the B isoform of Fru^M^ in the developing ovary did not inhibit TF formation and only induced weak esg-LacZ expression in the apical cap (Figure 5F). This indicates that the sex-specific function of *fru* in the gonad is regulated by Fru^Com^ rather than Fru^M^ and that Fru^Com^ has a stronger masculinizing effect than Fru^M^. Overall, we conclude that, while overexpression of Fru^B^ is sufficient to interfere with ovary development and partially masculinize somatic cells, it is not, by itself, sufficient to induce hub formation.

**Figure 5.**
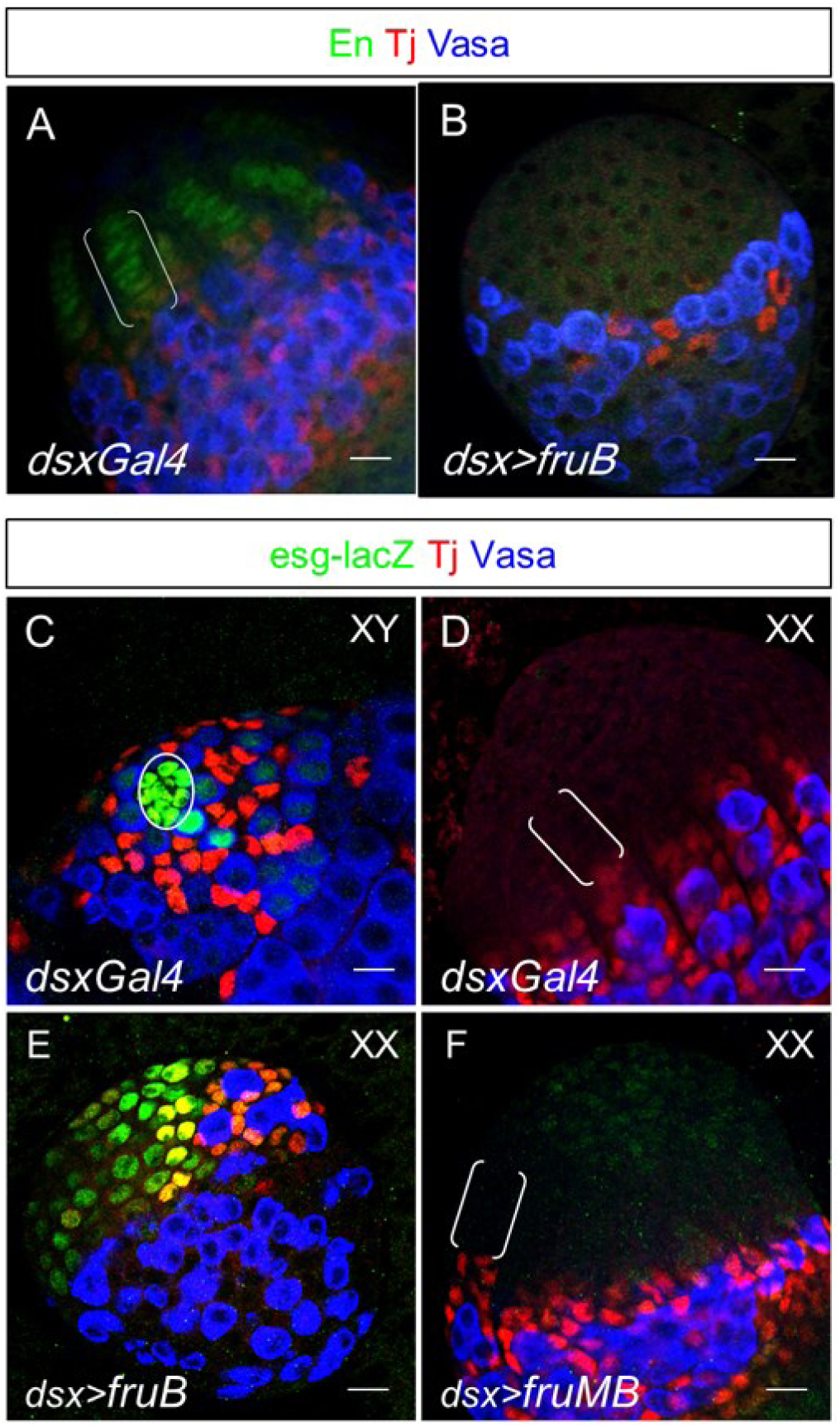
**Fru overexpression inhibits terminal filament development and masculinizes the female niche.** Ovaries of the white prepupal stage (WPP). Genotype as indicated. (C-F) WPP stage gonads. Genotype and chromosomal sex as indicated. Circle denotes the hub; brackets denote the TF. Scale bars represent 20 µm.

### An evolutionarily conserved Dsx binding site is required for normal expression from *fru* P4 promoter in hub cells

Previously, we have used a combination of whole-genome Dsx occupancy data, sequence searches for biochemically and genomically defined Dsx binding sites, and evolutionary conservation of these sites across sequenced Drosophila species, to identify likely Dsx targets in the genome [15]. This work indicated that *fru* was a candidate for direct regulation by Dsx, with the regions around the P3 and P4 promoters being particularly likely to contain Dsx-responsive elements (Figure S5). We identified a Dsx motif (DSX1) 6.3 kb upstream of P4 which is completely conserved across 21 *Drosophila* species and is a perfect match to the Dsx core binding motif (ACAATGT, [15, 28, 37]) and also matched surrounding nucleotides that may be important for Dsx binding [38] (Figure 6A and S6A). A transgenic reporter was created in which a 7.5 kb genomic sequence including DSX1 and the P4 promoter was placed upstream of a nuclear GFP reporter (WT reporter, Figure 6A). Transgenic flies carrying this construct (WT) expressed GFP in the hub, but not in the CySC or cyst cells, and expression was also not observed in the ovary in both late L3 and adult stages (Figure 6B and S6B, date not shown). Based on what we know about regulation of the few Dsx targets that have been studied, sex-specific expression in a given tissue requires both tissue-specific control elements and Dsx-responsive elements. Thus, it is not surprising that the WT *fru* reporter would be expressed in only a subset of Fru-expressing cells in the testis.

**Figure 6.**
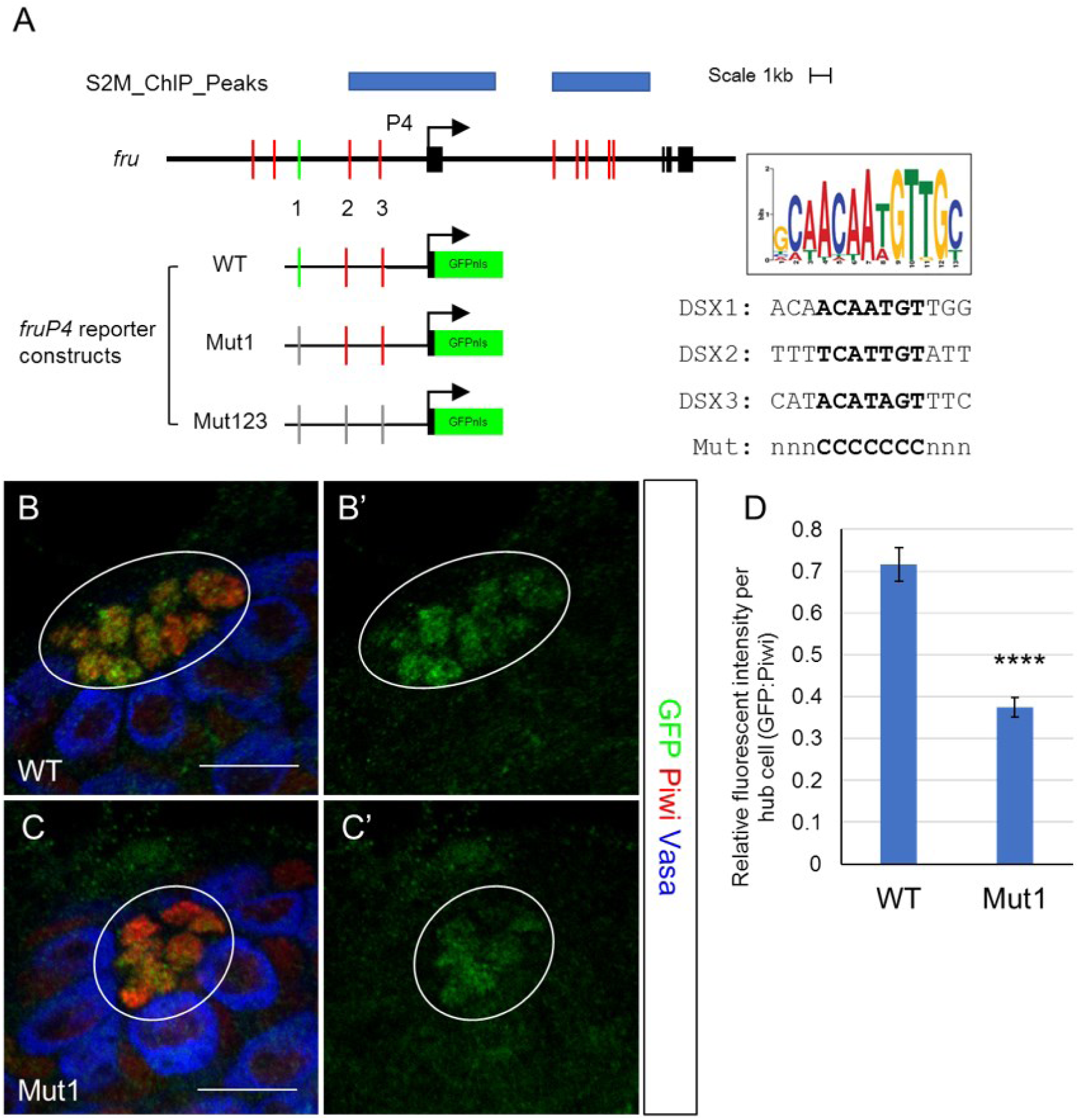
**A canonical Dsx binding site upstream of *fru* P4 is required to maintain normal P4 expression level in hub cells.** (A) Schematic diagram of the *fru* P4 enhancer-promoter constructs. Dsx occupancy (blue), and Dsx binding motifs (top 10% PWM, red and green bars) are shown. The Dsx consensus motif and the sequences of DSX1, DSX2, and DSX3 are shown. (B-C) Representative images of GFP expression levels in late L3 stage testes carrying WT and Mut1 constructs. Scale bars represent 20 µm. Circle denotes the hub. (D) Quantification of GFP relative fluorescent intensity per hub cell in WT and Mut1 testes. Data are presented as Mean ± SEM. WT, n=125; Mut1, n=115. Student’s t-test.

To test if DSX1 is essential for proper sex-specific expression of *fru*, we created the Mut1 reporter construct where the 7 core nucleotides of DSX1 are replaced by G-C base pairs. When GFP expression level in the hub was compared between transgenic flies containing WT and Mut1 constructs, we found that the GFP fluorescence intensity in hub cells of the Mut1 reporter was significantly decreased relative to the wild-type reporter (p<0.0001, student t-test, see Methods for quantification procedure) (Figure 6B-6D). However, we did not observe GFP expression in females which is expected if DsxF acts as a repressor of *fru* in the ovary (Figure S6C). Perhaps this is not surprising given the low level of Fru expression we observed in *dsx* mutants that formed female niche structures. Two other sites within the reporter transgene more weakly resemble the Dsx consensus motif, but are divergent in the 7-nucleotide core region (DSX2 and DSX3, Figure 6A). Mutation of these sites (Mut123) did not further decrease GFP expression in the hub or lead to GFP expression in the ovary (Figure S6D and S6E).

Collectively, these results support that *fru* is a direct target gene of Dsx. The conserved DSX1 motif is needed for robust expression in hub cells, but additional Dsx binding sites present in the *fru* locus, as well as additional tissue-specific elements, are likely needed to completely recapitulate sexually-dimorphic Fru expression in the gonad.

## DISCUSSION

Over the past decades, much effort has been focused on understanding the functions of *fru* in regulating sex-specific courtship behaviors, yet it remained unclear whether *fru* plays a role in regulating sexual dimorphism outside the nervous system. The work presented here demonstrates that Fru is expressed male-specifically in the gonad stem cell niche, and is required for CySC maintenance, cyst cell survival, and for the maintenance of the hub during larval development. Further, male-specific expression of Fru in the gonad is independent of the previously described mechanism of sex-specific alternative splicing by Tra, and is instead dependent on *dsx. fru* appears to be a direct target for transcriptional regulation by Dsx. This work provides evidence that *fru* regulates sex-specific development outside the nervous system and alters traditional thinking about the structure of the *Drosophila* sex determination pathway.

### *fru* function outside the nervous system

While it was previously reported that *fru* is expressed in tissues other than the nervous system, including in the gonad [13], a function for *fru* outside the nervous system was previously unknown. We find that Fru is expressed in the developing and adult testis in the hub, the CySC, and the early developing cyst cells. Importantly, we find that *fru* is important for the proper function of these cells.

Fru is not expressed at the time of hub formation during embryogenesis, but expression is initiated during the L2/L3 larval stage. This correlates with a time period when the hub must be maintained and resist transforming into female niche structures; in *dsx* mutants, all gonads in XX and XY animals develop hubs, but half of each transform into terminal filament cells and cap cells (Camara *et al.*, submitted). *fru* is not required for initial hub formation, consistent with it not being expressed at that time. *fru* is also not, by itself, required for hub maintenance under the conditions that we have been able to assay (prior to the pupal lethality of *fru* null mutant animals). However, under conditions where hub maintenance is compromised by loss of *dsx* function, *fru* clearly plays a role in influencing whether a gonad will retain a hub, or transform into TF. Fru expression in *dsx* mutant gonads correlates with whether they will form male or female niche structures (Figure 2H), and removing even a single allele of *fru* is sufficient to induce more hubs to transform into TFs (Figure 3). The mechanisms that influence the variable levels of Fru expression in the absence of *dsx* remain unknown. Finally, ectopic expression of Fru in females is sufficient to inhibit TF formation and partially masculinize the gonad (Figure 5B and 5E), but does not induce hub formation. Thus, we propose that *fru* is one factor acting downstream of *dsx* in the maintenance of the male gonad stem cell niche, but that it acts in combination with other factors that also regulate this process.

We also demonstrated that *fru* is required for CySC maintenance and also for the survival of differentiating cyst cells and their germline cysts. Loss of *fru* from the CySC lineage led to rapid loss of these CySC from the testis niche (Figure 4A). Since we did not observe precocious differentiation of these cells or an increase in their apoptosis (Figure 4B and 4C), these mechanisms do not appear to contribute to CySC loss. One possibility is that *fru* is needed for CySCs to have normal expression of adhesion proteins and compete with other stem cells for niche occupancy. It has been shown that *fru* regulates the Slit-robo pathway and *robo1* is a direct target of *fru* in the CNS [9, 39]. Interestingly, the Slit-Robo pathway also functions in the CySCs to modulate E-cadherin levels and control the ability of CySCs to compete for occupancy in the niche [40]. Therefore, *fru* may use similar mechanisms to maintain CySCs attaching to the hub.

*fru* also influences survival in the differentiating cyst cells, as we observed an increase in cell death in these cells in *fru* mutants. Several reports have demonstrated that *fru* represses programmed cell death in the nervous system [7, 8, 41]. It was further indicated that the cell death gene *reaper* is a putative target of Fru [32]. *fru* is also required for survival of trans-amplifying spermatogonial cysts, and this is likely indirectly due to *fru’s* role in cyst cell survival.

In summary, *fru* function is clearly important for male niche maintenance and the function of the CySC and their differentiating progeny. This provides clear evidence that *fru* regulates sex-specific development in tissues other than the nervous system. Whether other tissues are also regulated by *fru* remains to be determined.

Interestingly, our data also support the idea that different isoforms of Fru also can have very different functions. In the CNS, Fru^A^, Fru^B^, and Fru^C^ have mostly overlapping expression, but play distinct roles: Fru^B^ is required for all steps of courtship behaviors; FruC controls specific steps; and Fru^A^ is dispensable for courtship behaviors [42, 43]. In the gonad, we also detected all C-terminal isoforms of *fru* at the mRNA level (Figure S3B). We show that Fru^B^ and Fru^C^, but not Fru^A^, are required for *fru* functions in hub maintenance (Figure 3B). Additionally, using clonal analysis, we show that Fru^B^ plays a more important role than Fru^C^ in the maintenance of the CySC fate (Figure 4A). Interestingly, we see a dramatic loss of all Fru^Com^ immunoreactivity in the testis in the *fru*^*B*^ isoform mutant, even though RT PCR indicated that multiple Fru isoforms are expressed in the testis. This suggests that either Fru^B^ is the predominant isoform expressed in the testis, or that loss of Fru^B^ function results in loss of other Fru isoforms, possibly through increased degradation (implying protein-protein interaction) or reduced *fru* transcription (implying autoregulation of the *fru* locus by Fru^B^).

In addition, proteins generated from the P1 promoter (Fru^M^) differ from other isoforms by the presence of a 101 sequence N-terminal extension. It has been shown that the functions of *fru* cannot be replaced by Fru^M^ during embryonic CNS development [34], suggesting that this N-terminal extension alters Fru activity. We find that ectopic expression of Fru^MB^, which contains the N-terminal extension, has a much less potent effect in masculinizing the female gonad than Fru^B^ (Figure 5E and 5F). Thus, in both the nervous system and the gonad, the Fru^M^ proteins appear to have different effects than the Fru^Com^ proteins, though how the N-terminal extension of Fru^M^ modulates its activity is unknown.

### A change in our view of the sex determination pathway

Previously, it was thought that the only mechanism by which sex-specific functions of *fru* were regulated was through Tra-dependent alternative splicing of the P1 transcripts. *fru* null alleles are lethal in both sexes and Fru protein encoded by non-P1 *fru* transcripts was thought to be sex-nonspecific and to not contribute to sex determination. Thus, *fru* and *dsx* were thought to be parallel branches of the sex determination pathway, each independently regulated by Tra. Here we demonstrate that *fru* can also be regulated in a manner independent of *tra* and dependent on *dsx*, and provide evidence that *fru* is a direct target for transcriptional regulation by Dsx (Figure 7). First, Fru expression in the testis is independent of the P1 transcript that is regulated by Tra. A P1 reporter is not expressed in the testis and a mutation that prevents Fru^M^ expression from P1 does not affect Fru immunoreactivity in the testis (Figure S1G and S1H). Second, in animals that simultaneously express the female form of *tra* (Tra on) and the male form of Dsx (XX; *dsx*^*D*^/*dsx*^*0*^), Fru is expressed in the male mode in the testis, demonstrating that it is regulated by *dsx*. Finally, an evolutionarily conserved Dsx consensus binding site upstream of the *fru* P4 promoter is required for proper expression levels of a *fru* P4 reporter in the testis. Together, these data demonstrate a novel mode for *fru* regulation by the sex determination pathway, where sex-specific functions of *fru* are regulated by *dsx*. This also means that the large number of *fru* transcripts that do not arise from the P1 promoter can be expressed in a sex-specific manner to contribute to sexual dimorphism.

**Figure 7.**
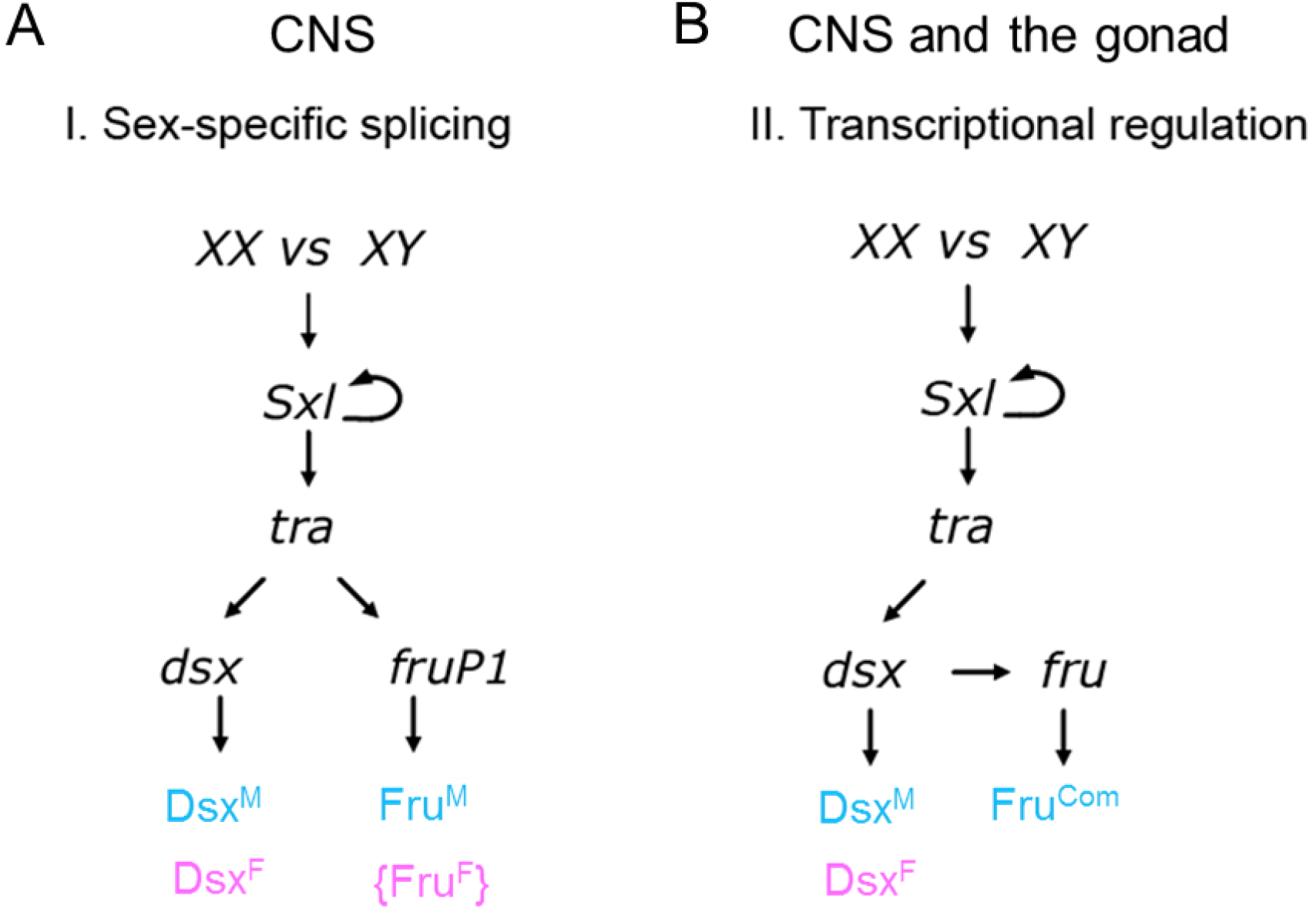
**Proposed model of the *Drosophila* sex determination pathway.** (A) The canonical sex determination pathway controls male-specific Fru^M^ expression in the CNS through sex-specific splicing of *fru* P1 transcripts. (B)The non-canonical sex determination pathway controls male-specific Fru^Com^ expression in the CNS and the gonad through transcriptional regulation by Dsx^M^ and Dsx^F^. Fru^M^ functions in parallel with Dsx to control sexual dimorphism in courtship behavior. Fru^Com^ functions downstream of Dsx to regulate sexual dimorphism of the gonad stem cell niche and potentially the nervous system.

The male and female forms of Dsx contain the same DNA binding domain and are thought to regulate the same target genes, but to have opposite effects on gene expression. Prior to this study, the two documented Dsx targets (*Yolk proteins 1* and *2* and *bric-a-brac*), along with other proposed targets, were all expressed at higher levels in females than males [28–30, 44]. Thus, for these targets, DsxF acts as an activator and DsxM acts as a repressor. Interestingly, *fru* is the first Dsx target gene that is expressed in a male-biased, rather than female-biased, manner. Thus, for direct regulation of *fru*, DsxM would activate expression while DsxF represses. Mechanistically for Dsx, this implies that the male and female isoforms are not dedicated repressors and activators, respectively, but may be able to switch their mode of regulation in a tissue-specific or target-specific manner. The mouse DMRT1 has also been shown to regulate gene expression both as transcriptional activator and repressor [45]. Thus, it is quite possible that bifunctional transcriptional regulation is a conserved characteristic of DMRTs.

Interestingly, we find evidence that *fru* is also regulated in the brain in a manner that is dependent on *dsx* and independent of *tra* (Figure 2D-2G). Thus, *dsx* regulation of *fru* occurs in the nervous system as well, where it co-exists with direct regulation of *fru* alternative splicing by Tra. It was originally thought that alternative splicing of the *fru* P1 transcript was essential for male courtship behavior [23]. However, more recently it was found that simply altering the courtship assay, by allowing *fru* P1-mutant males to be stimulated by other flies prior to testing, revealed that these animals could, in fact, exhibit male courtship behavior [5]. Interestingly, the courtship behavior exhibited by these males was dependent on *dsx*. We propose that *fru* might still be essential for male courtship in these *fru* P1-mutants, but that now sex-specific *fru* expression is dependent on transcriptional regulation by *dsx.*

### Evolution of the sex determination pathway

If sex-specific *fru* function can be regulated both through alternative splicing by Tra and through transcriptional regulation by Dsx, it raises the question of what is the relationship between these two modes of regulation. We propose that regulation of *fru* by Dsx is the more ancient version of the sex determination pathway and that additional regulation of *fru* by Tra evolved subsequently, through the acquisition of cis-regulatory elements in the *fru* P1 intron-exon splicing junction. This model is supported by studies of *fru* gene structures in distantly related Dipteran species and species of other insect orders. Studies of different insect species illustrate the considerable variability in the organization of *cis* sequences controlling *fru* alternative splicing [46]. Further, in some insects, no evidence for alternative splicing of *fru* at all has been found, yet *fru* still plays an important role in males to control courtship behaviors [47–49]. And in two Hawaiian picture-winged groups of subgenus *Drosophila*, the *fru* orthologues lack the P1 promoter, and non-P1 *fru* transcripts exhibit male-specific expression [50, 51], similar to what we propose for non-P1 *fru* transcripts in *D. melanogaster*. Thus, it appears that regulation of *fru* by *dsx* may be the evolutionarily more ancient mechanism for sex-specific control of *fru*, while Tra-dependent splicing of P1 transcripts is a more recent adaptation. More broadly, *tra* is not conserved in the sex determination pathway in the majority of animal groups, while homologs of Dsx, the DMRTs, are virtually universal in animal sex determination. Thus, if Fru-related transcription factors are involved in the creation of sexual dimorphism in the body or the brain in other animals, they cannot be regulated by Tra and may be regulated by DMRTs.

## METHODS

### Fly Stocks

The following stocks were used: *fru*^*W24*^ (S. Goodwin), *fru*^*Sat15*^ (S. Goodwin), *fru*^Δ*A*^ (S. Goodwin), *fru*^Δ*B*^ (S. Goodwin), *fru*^Δ*C*^ (S. Goodwin), *dsx*^*D*^, *Df(3R)dsx*^*3*^, *dsx*^*1*^, *dsx*^*Gal4*Δ*2*^ (B. Baker), *dsx-Gal4* (S. Goodwin), <u>*UAS-fruMB*</u> (S. Goodwin), *UAS-fruB* (S. Goodwin), *c587-Gal4* (T. Xie), *tj-Gal4* (D. Godt), *tub-Gal80*^*ts*^, *esg*^*M5-4*^ (S. DiNardo), *y^1^ v^1^; P{TRiP.JF01182}attP2* (*UAS-fru*^*Com*^-*RNAi*), *yw, hs-FLP,UAS-mCD8:GFP;tub-Gal4,neoFRT82B,tub-Gal80*, *hs-FLP, tub-Gal4, UAS-GFP.Myc.nls, yw; neoFRT82B, tub-Gal80*, *FRT82B*, *FRT82B, fru*^*Sat15*^, *FRT82B, fru*^Δ*B*^, *FRT82B, fru*^Δ*C*^ and *w*^*1118*^ as a control. All flies were raised at 25 ºC unless otherwise stated.

### Immunohistochemistry

Adult testes were dissected in PBS and fixed at room temperature for 15 minutes in 4.5% formaldehyde in PBS containing 0.1% Triton X-100 (PBTx). Adult ovaries, *dsx* mutant adult gonads, and larval gonads were dissected in PBS followed by a 10-minute fixation at room temperature in 6% formaldehyde in PBTx. Immunostaining was performed as previously described (Gonczy *et al*., 1997), and samples were mounted in 2.5% DABCO. Adult CNS The following primary antibodies were used: rat anti-Fru^Com^ at 1:300 (S. Goodwin); guinea pig anti-Traffic-jam (D. Godt) at 1:10,000; mouse anti-Arm N2 7A1 (DSHB, E. Wieschaus) at 1:100; chicken anti-Vasa (K. Howard) at 1:10,000; mouse anti-Fas-3 7G10 (DHSB, C. Goodman) at 1:30; mouse anti-Eya 10H6 (DSHB, S. Benzer/N.M. Bonini) at 1:25; mouse anti-Engrailed 4D9 (DSHB, C. Goodman) at 1:2; rat anti-DN-Cad DN-EX#8 (DHSB, T. Uemura) at 1:20; rabbit anti-GFP (Abcam) at 1:2000; rabbit anti-Vasa (R. Lehmann) at 1:10,000; rabbit anti-Sox100B (S. Russell) at 1:1,000; rabbit anti-β-Gal (Cappel) at 1:10,000; rabbit anti-Zfh1 (R. Lehman) at 1:5,000. Secondary Alexa 488, 546 and 633 antibodies were used at 1:500 (Invitrogen, Carlsbad, CA). All immunohistochemistry samples were imaged on a Zeiss LSM 700 confocal microscope.

### Developmental staging

To obtain first and second instar larvae, flies were transferred to a cage to allow egg-laying on an apple juice plate for 4 hours and were then removed. The apple juice plates were left at 25°C. Larvae were collected at desired developmental stages (36 h for mid first instar, 72 h for late second instar). Immobile third instar larvae were collected from vials as late third instar larvae. Larvae with inverted spiracles and harden carcass were collected from vials as white prepupae.

### Genotyping and sex identification of dsx mutants

Balancer chromosomes containing a *P{Kr-GFP}* transgene were used to distinguish transheterozygous *dsx* or *fru* mutant larvae from heterozygous siblings. Sex chromosome genotype of *dsx* null mutants was identified using a *P{Msl-3-GFP}* (J. Sedat) transgene, or Y chromosome marked with *Bs* (*Dp(1;Y)B*^*S*^). XX *dsx*^*D*^/+ and *dsx*^*D*^/*dsx*^*0*^ mutants were distinguished from XY siblings by abnormal gonad morphology.

### Quantification of niche identity

Adult flies less than 2 days old were dissected and stained with antibodies against DN-Cad, Fas-3, and Vasa, and cell nuclei were visualized via DAPI staining. Z-stack images were taken with a Zeiss LSM 700 confocal microscope with a 40x objective. The hub was defined as a compact cluster of DAPI bright somatic cells that co-expressed N-cad and Fas-3 and were surrounded by a rosette of Vasa-positive germ cells. TFs were determined by ladder-shaped N-Cad staining around stacks of disc-shaped somatic nuclei indicated by DAPI staining. Gonad was defined as no niche when neither TFs nor a hub was identified.

### Clonal analysis

Flies of the following genotype were used for MARCM: *hs-FLP, UAS-mCD8:GFP/Y; tub-Gal4, FRT82B, tub-Gal80/FRT82B* (control 1); *hs-FLP, UAS-mCD8:GFP/Y; tub-Gal4, FRT82B, tub-Gal80/FRT82B, ry* (control 2); *hs-FLP, UAS-mCD8:GFP/Y; tub-Gal4, FRT82B, tub-Gal80/FRT82B, fru*^*Sat15*^; *hs-FLP, UAS-mCD8:GFP/Y; tub-Gal4, FRT82B, tub-Gal80/FRT82B, fru*^Δ*B*^; *hs-FLP, UAS-mCD8:GFP/Y; tub-Gal4, FRT82B, tub-Gal80/FRT82B, fru*^Δ*C*^. Newly eclosed adult males (0-2 day old) were collected at 25 ºC prior to heat shock. Flies were heat-shocked at 37 ºC for 1 hr and returned to 25 ºC and raised in fresh vials with yeast paste. Control and mutant clones were analyzed at the indicated time points post clonal induction. CySC clones were counted as GFP-marked Zfh-1- or Tj-positive cells within one cell diameter to the hub and directly contacting the hub with cytoplasmic extension as indicated by mCD8:GFP. Rest of the GFP marked Zfh-1- or Tj-positive cells are considered as cyst cell clones.

### RT-PCR

50 pairs of late 3^rd^ instar larval gonads were dissected into ice-cold PBS and cDNA was prepared following manufacturers’ protocols (Zymo Research Quick-RNA Miniprep Kit and Invitrogen Superscript III Kit). PCR was performed on cDNA using the following intron-spanning primer pairs (given in the 5’-3’ orientation):

> RP48-F - CCGCTTCAAGGGACAGTATCTG
>
> RP48-R - ATCTCGCCGCAGTAAACGC
>
> TJ-F - ACCAGTGGCACATGGACGAA
>
> TJ-R - CGCTCCCGAAGATGTGTTCA
>
> Actin5C-F - TAATCCAGAGACACCAAACC
>
> Actin5C-F – CAGCAACTTCTTCGTCACAC
>
> Fru-P1-F - CGGAAAAGGGCGTATGGATTG
>
> Fru-P1-R - TGTGCCAGTCAGCCTCTG
>
> Fru-P2-F - AGCACGCCGGTCAAATTTG
>
> Fru-P2-R - TCGCTCGGTTTTAGTTTCCCA
>
> Fru-P3-F - GCACGTTCTCAGTTTGGAATTC
>
> Fru-P3-R - CAACGAAAACCGTGAACTGTG
>
> Fru-P4-F - GAATTGCTGGTCCATCGCTC
>
> Fru-P4-R - GCAACTGAACCCAACTGTACC
>
> Fru-Com-F - ATTACTCGGCCCACGTCC
>
> Fru-Com-R - CTGCCCATGTTTCTCAAGACG
>
> Fru-A-F - GCTGGACCAGACGGACAATA
>
> Fru-A-R - GTCGTGCTCCCGATGATTT
>
> Fru-B-F - same as Fru-A-F
>
> Fru-B-R - CAACGGTGCAGGTTGCAG
>
> Fru-C-F - same as Fru-A-F
>
> Fru-C-R - GACAGGTGCATCCCGAAAG
>
> Fru-D-F - CCAGATTACTTGCCGGTGAA
>
> Fru-D-R - GCTCTTCAACTGAGCCTCCA

Each primer pair was validated for efficacy using whole fly cDNA from adult male (data not shown).

### Fru reporter constructs and transgenes

To generate the WT fru P4 enhancer-promoter reporter construct, a 7.5 kb genomic sequence from *fru* genomic clone BACRP98-2G21(BACPAC Resources Center) was amplified with the following primers (given in the 5’ to 3’ orientation) and cloned into the pJR16 vector (R. Johnston) between the BamHI and PstI site.

> Fru-P4-8K-WT-F - CGGGATCCGCAACCCGTCCGTATC
>
> Fru-P4-8K-WT-R - CAACTGCAGTGTGGGTATGGGCAAATTGA

Site-directed mutagenesis of DSX sites was performed according to the manufacturer’s protocol (New England Biolabs Q5 Site-Directed Mutagenesis Kit). The following primer sets were used:

> DSX1mut-F - GGGTGTGTTAATTTGCCAGG
>
> DSX1mut-R - CCCCTGGCTCATTAACAGACCAAT
>
> DSX2mut-F - GGGATTTATTGCACAGGTTG
>
> DSX2mut-R - CCCCAAATGTTAGAAAACCAAGCATTTTT
>
> DSX3mut-F - GGGTTCTGTAATAGATAATTCAGTTC
>
> DSX3mut-R - CCCCATGAGTAACTTCTGTGC

Transgenic flies were generated via PhiC31 integrase-mediated transgenesis. The constructs were integrated into the same genomic location (attP40 on Chromosome II).

### Imaging and quantification of GFP expression in the hub

Z-stack images of the hub were taken under the same setting on a Zeiss LSM 700 confocal microscope with a 63x objective. Quantification of GFP fluorescent intensity was performed in Fiji software (ImageJ, NIH). For each gonad, five random hub cells were sampled, and a 16-cell-stage germ cell was sampled as background. A circle of the same size was drawn as the sample area. Average fluorescence intensity of GFP and Piwi was acquired. The relative fluorescent intensity was measured as (GFP[hub]-GFP[background])/(Piwi[hub]-Piwi[background]).

## ACKNOWLEDGMENTS

We thank Stephen Goodwin for generously sharing *fruitless* reagents. We also thank the fly community, the Bloomington Stock Center, the Flybase, the BACPAC Resources Center and the Developmental Studies Hybridoma Bank for stocks, reagents and information. We thank the Johns Hopkins Community for helpful discussion. C.P. is supported by the NSF Graduate Research Fellowship Program (DGE1746891). This work was supported by a grant to M.V.D. from the National Institute of General Medical Sciences (GM113001).

## FIGURE LEGEND

**Supp Figure 1.**
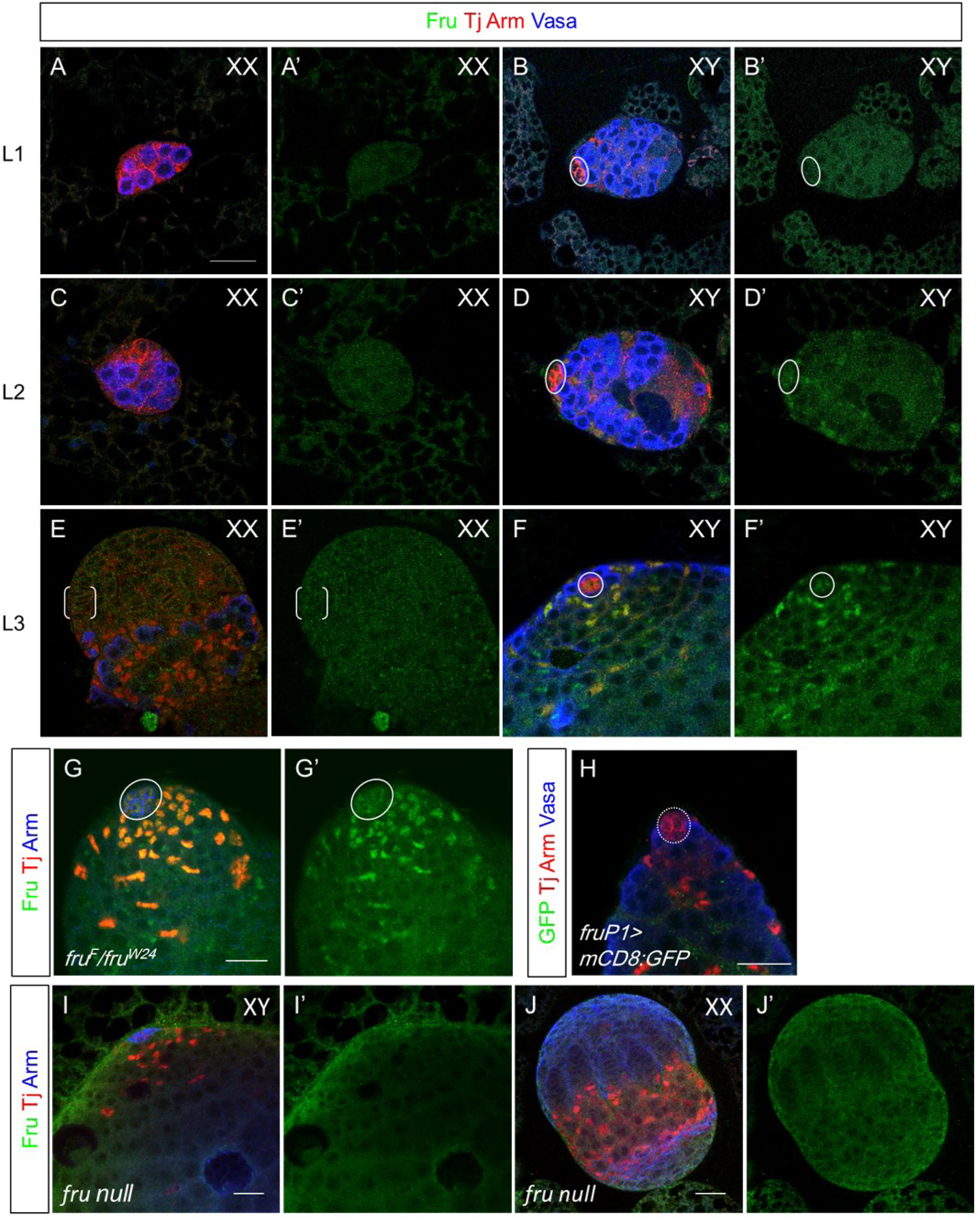
(A-B) Wildtype L1 stage female and male gonads with no Fru expression. (C-D) Wildtype L2 stage gonads showing weak Fru expression in the hub cells and early CySC lineage of the testis. (E-F) Wildtype late L3 stage gonads showing robust Fru expression in the male GSC niche and no Fru expression in the female GSC niche (G) A representative *fru*^*F*^/*fru*^*W24*^ adult testis showing normal Fru expression in the niche. (H)A representative *fru*^*P1Gal4*^>*mCD8:GFP* testis showing no GFP expression in the niche. (I-J) Late L3 stage male (I) and female (J) gonads showing the removal of the Fru^Com^ immunoreactivity in the GSC niche by *fru*^*Sat15*^/*fru*^*W24*^. Scale bars represent 20 µm. Circle denotes the hub; brackets denote the TF.

**Supp Figure 2.**
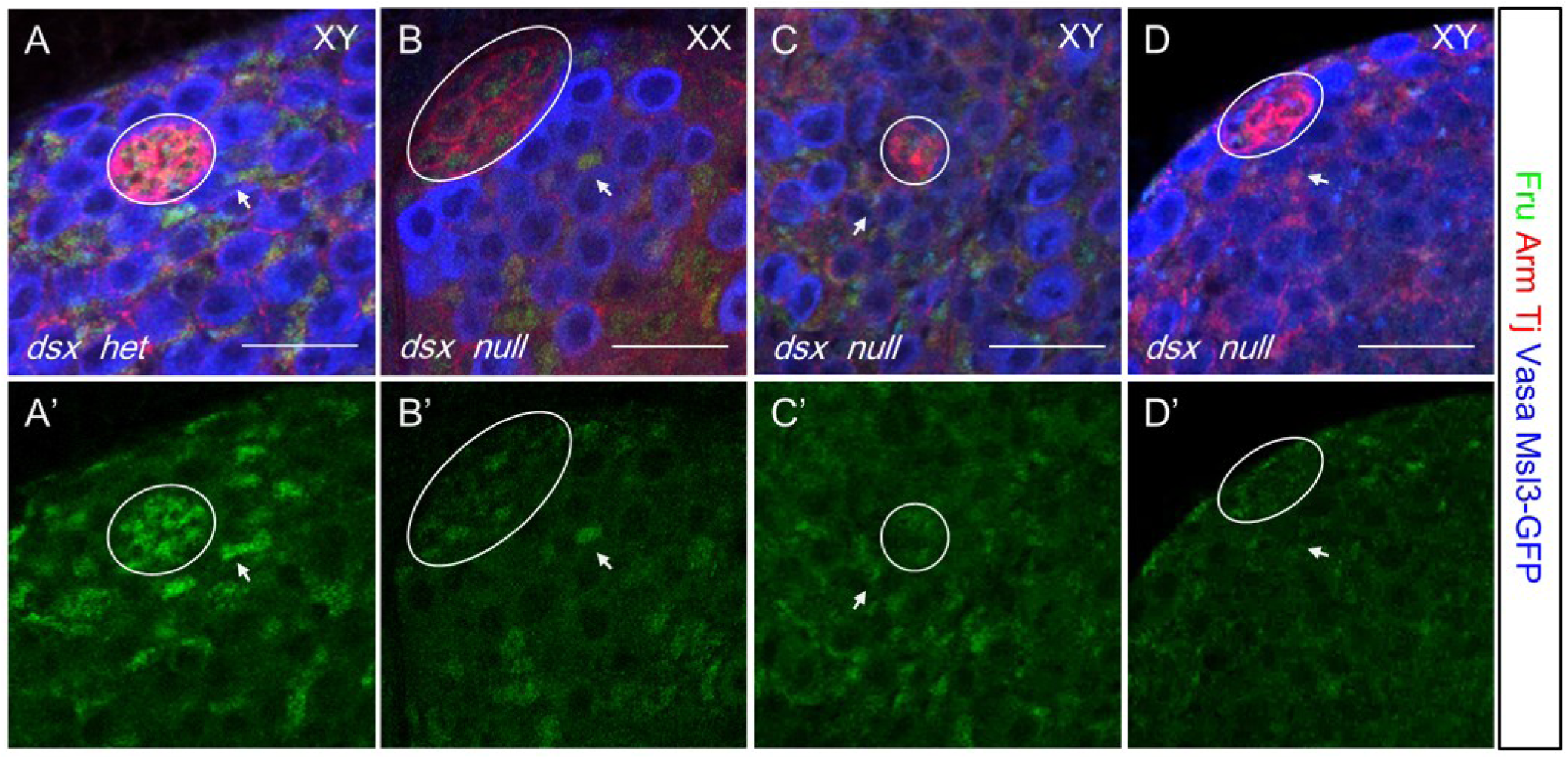
(A) A representative XY *dsx* heterozygote GSC niche showing wild-type level of Fru expression. (B) A representative XX *dsx*^*3*^/*dsx*^*1*^ gonad with the male niche identity showing Fru expression in the hub cells and early CySC lineage at a reduced level. (C-D) representative images showing XY *dsx*^*3*^/*dsx*^*1*^ gonads with a hub have variable Fru expression levels. All images represent late L3 stage gonads. Scale bars represent 20 µm. Circle denotes the hub; brackets denote the TF; arrows denote CySCs.

**Supp Figure 3.**
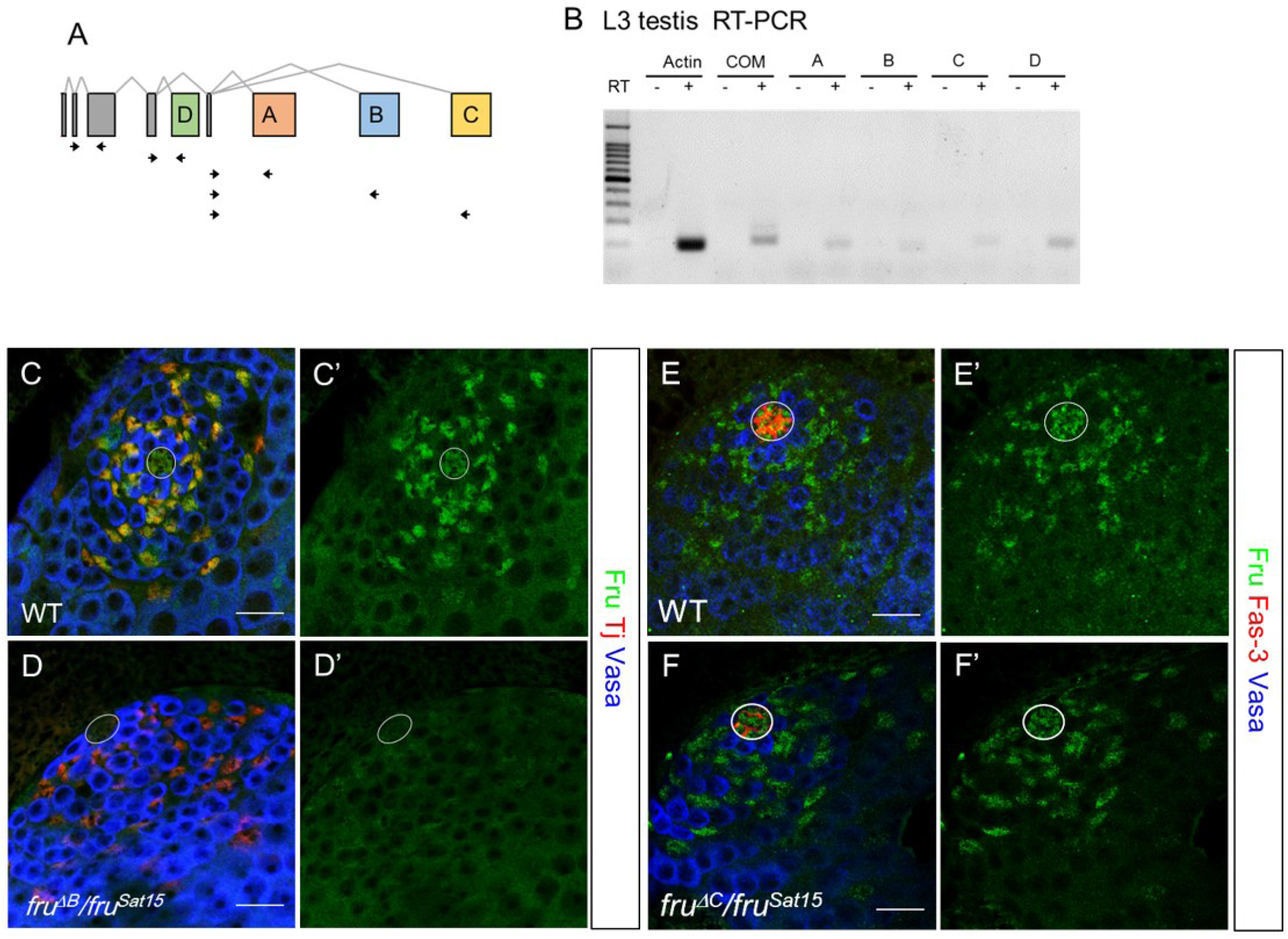
(A) Diagram showing sex-nonspecific splicing of the C-terminal isoforms. A, B, C, and D isoform each contains a distinct zinc-finger domain. Intron-spanning primer sets used to probe the isoforms are as indicated. (B) RT-PCR result using *actin*, *fruCOM*, or isoform-specific *fru* primers. (C-D) Wildtype (C) and *fru^ΔB^/fru*^*Sat15*^ (D) late L3 stage testes. (E-F) Wildtype (E) and *fru*^Δ*C*^/*fru*^*Sat15*^ (F) late L3 stage testes. Circle: the hub; arrow: residual Fru expression in the cyst cell. Scale bars represent 20 µm. Circle denotes the hub.

**Supp Figure 4.**
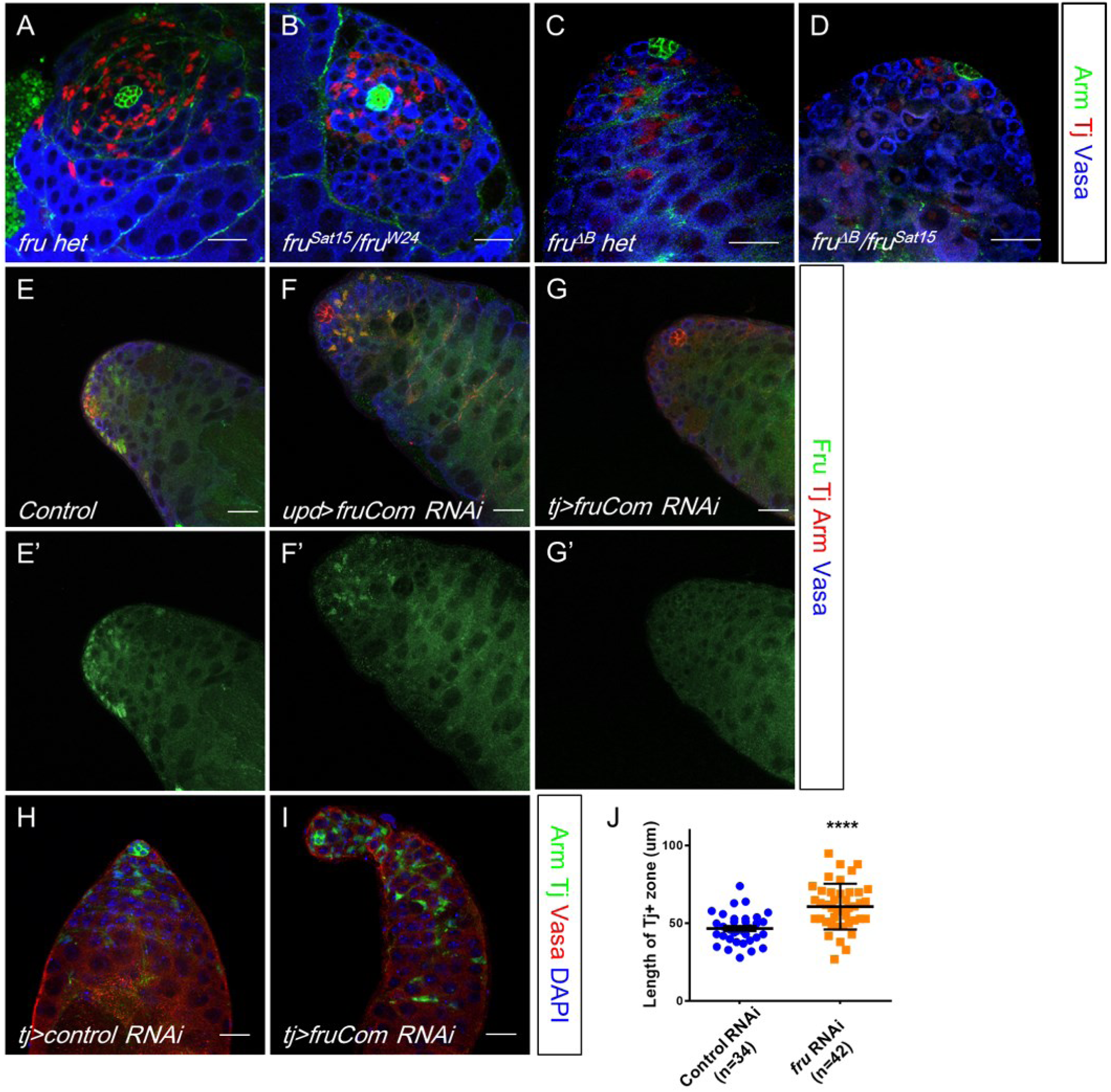
(A-B) White prepupal stage *fru* het (A) and *fru* null (B) testes. (C-D) Representative images of *fru* het (C) and *fru*^Δ*B*^ mutant (D) testes 3 days after puparium formation. (E-G) 1-week old testis with *UAS-fruCom RNAi* (E) alone, or expressing *fruCom RNAi* in the hub with *upd-Gal4* (F) or expressing *fruCon RNAi* in the hub and early CySC lineage with *tj-Gal4* (G). (H-I) 2-week old testes expressing *GFP RNAi* (H) or *fruCom RNAi* (I) with *tj-Gal4*. (J) Quantification of the length of TJ+ zone (20 µm) in control and *fruCom RNAi* testes. Mean ± SD, Student’s t-test. Scale bars represent 20 µm. Circle denotes the hub.

**Supp Figure 5.**
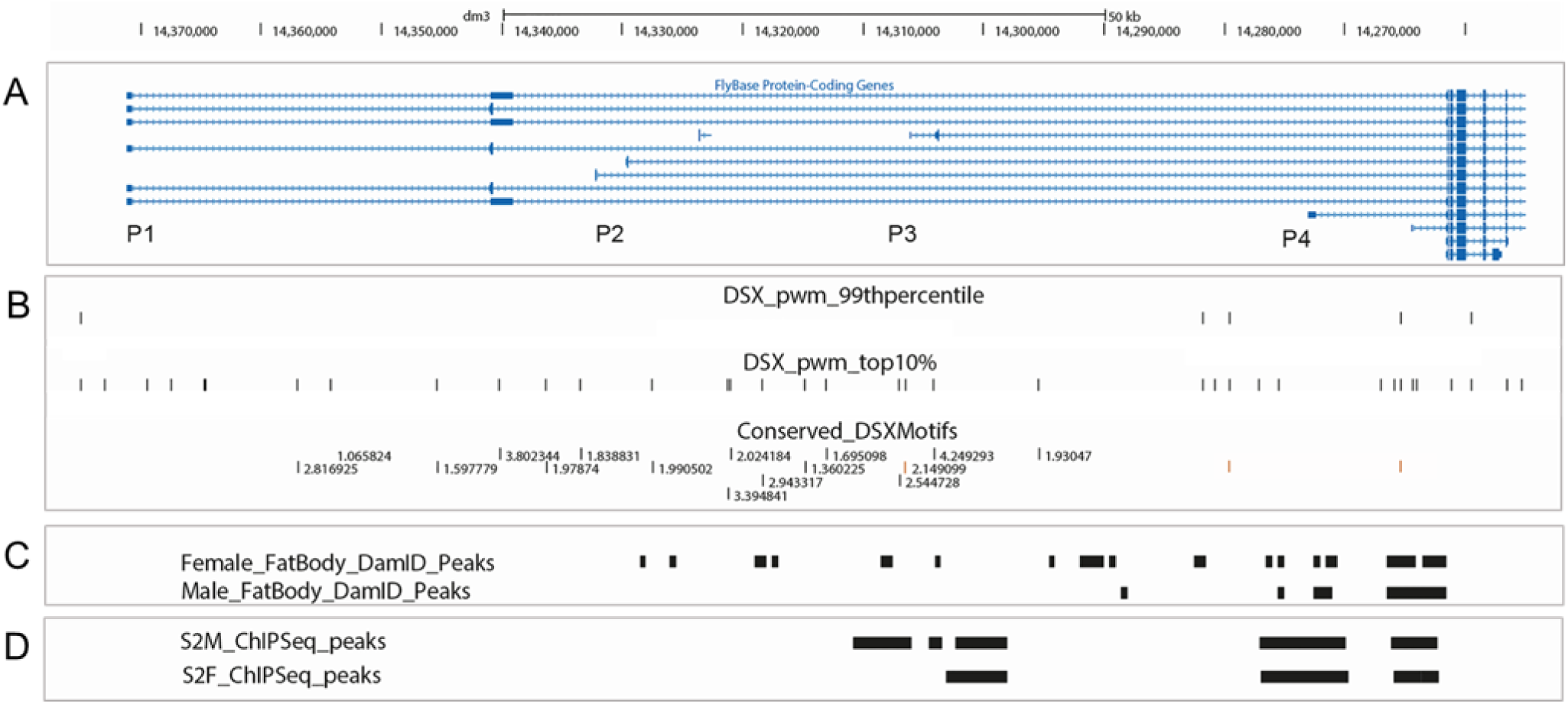
(A-D) The *fru* promoter region is shown to scale with transcripts generated from P1-P4 are labeled. Putative Dsx binding motifs shown as top 1% PWM, top 10% PWM and evolutionarily conserved Dsx motifs (B). Female and male fate body Dsx-DamID (C) and S2 cells DsxM and DsxF ChIP-Seq(D) peaks.

**Supp Figure 6.**
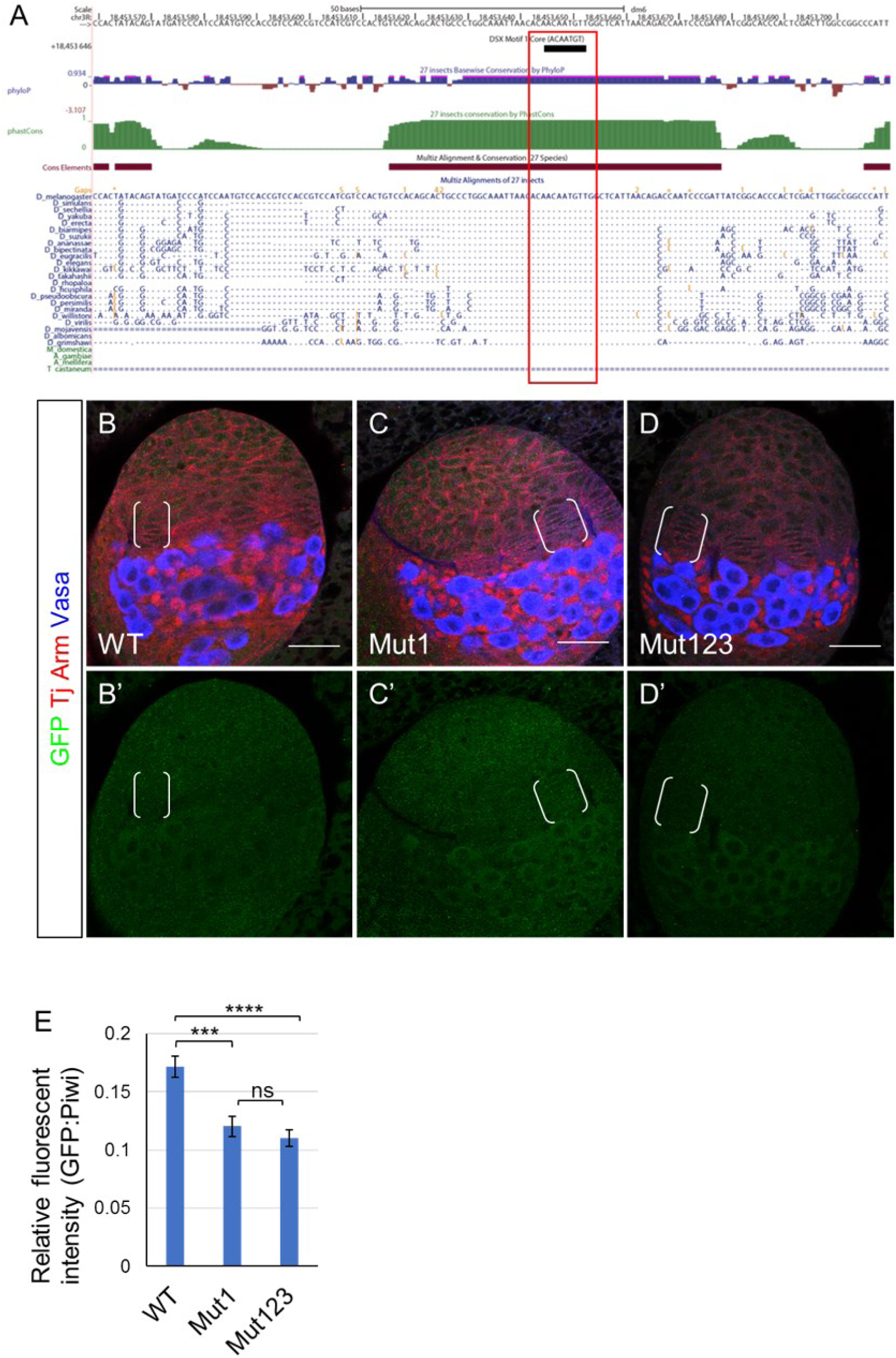
(A) Evolutionary conservation analyses of DSX1 using comparative genomics tracks of the UCSC Genome Browser. Sequence alignment among *Drosophila* species is shown with same nucleotides abbreviated as dots. (B-D) GFP expression of P4 WT (B), Mut1 (C) and Mu123 (D) constructs in late L3 stage ovaries. Scale bars represent 20 µm. Brackets denote TFs. (E) Comparison of relative GFP fluorescent intensity per hub cells (standardized by Piwi expression) in WT, Mut1 and Mut 123 constructs. Bars represent Mean±SEM. Sample size: WT, n=50; Mut1, n=35; Mut123, n=40. Student’s t-test.

**Supp Table 1.** Quantification of niche sex identity in *dsx*^*3*^/*dsx*^*1*^ and *XX*; *dsx*^*D*^/+ adult gonads.

| Genotype | Sample size | Hub% (AVG±SD) | TF% (AVG±SD) | No niche% (AVG±SD) |
| --- | --- | --- | --- | --- |
| *dsx<sup>3</sup>/dsx<sup>1</sup>*<br>XY | 32 | 40.6±15.0 | 43.8±0.8 | 15.6±14.1 |
| *dsx<sup>3</sup>, fru<sup>Sat15</sup>/dsx<sup>1</sup>*<br>XY | 50 | 8.0±3.6 | 64.0±12.7 | 28.0±16.4 |
| *dsx<sup>3</sup>, fru<sup>Sat15</sup>/dsx<sup>1</sup>*<br>XX | 44 | 13.6±4.3 | 72.7±7.1 | 13.6±5.0 |
| XX *w<sup>1118</sup>/dsx<sup>D</sup>* | 110 | 26.4±3.9 | 62.7±3.2 | 10.9±3.3 |
| XX *fru<sup>Sat15</sup>/dsx<sup>D</sup>* | 256 | 4.7±4.1 | 87.5±6.1 | 7.8±5.5 |
| XX *fru<sup>ΔB</sup>/dsx<sup>D</sup>* | 179 | 1.7±2.4 | 97.8±2.3 | 0.6±1.5 |
| XX *fru<sup>ΔC</sup>/dsx<sup>D</sup>* | 163 | 3.1±6.9 | 92.0±7.2 | 4.9±6.6 |
| XX *fru<sup>ΔA</sup>/dsx<sup>D</sup>* | 63 | 36.5±10.5 | 49.2±12.1 | 14.3±4.0 |

**Supp Table 2.**
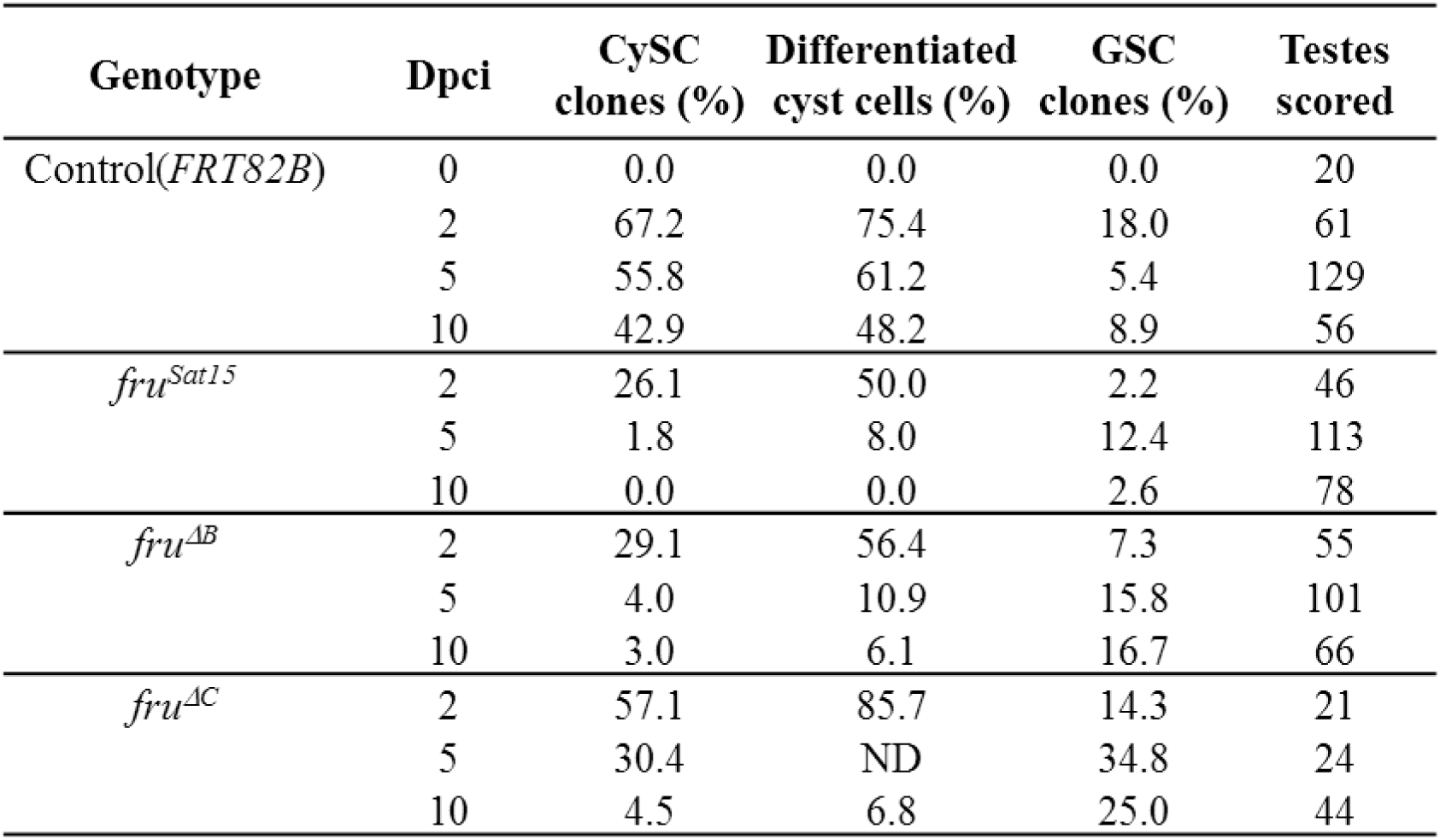
Quantification of control and *fru* clones.

